# Enhancer variants associated with Alzheimer’s disease affect gene expression via chromatin looping

**DOI:** 10.1101/426312

**Authors:** Masataka Kikuchi, Norikazu Hara, Mai Hasegawa, Akinori Miyashita, Ryozo Kuwano, Takeshi Ikeuchi, Akihiro Nakaya

## Abstract

**Background:** Genome-wide association studies (GWASs) have identified single-nucleotide polymorphisms (SNPs) that may be genetic factors underlying Alzheimer’s disease (AD). However, how these AD-associated SNPs (AD SNPs) contribute to the pathogenesis of this disease is poorly understood because most of them are located in non-coding regions, such as introns and intergenic regions. Previous studies reported that some disease-associated SNPs affect regulatory elements including enhancers. We hypothesized that non-coding AD SNPs are located in enhancers and affect gene expression levels via chromatin loops.

**Results:** We examined enhancer locations that were predicted in 127 human tissues or cell types, including ten brain tissues, and identified chromatin-chromatin interactions by Hi-C experiments. We report the following findings: (1) nearly 30% of non-coding AD SNPs are located in enhancers; (2) expression quantitative trait locus (eQTL) genes affected by non-coding AD SNPs within enhancers are associated with amyloid beta clearance, synaptic transmission, and immune responses; (3) 95% of the AD SNPs located in enhancers co-localize with their eQTL genes in topologically associating domains suggesting that regulation may occur through chromatin higher-order structures; (4) rs1476679 spatially contacts the promoters of eQTL genes via CTCF-CTCF interactions; (5) the effect of other AD SNPs such as rs7364180 is likely to be, at least in part, indirect through regulation of transcription factors that in turn regulate AD associated genes.

**Conclusion:** Our results suggest that non-coding AD SNPs may affect the function of enhancers thereby influencing the expression levels of surrounding or distant genes via chromatin loops. This result may explain how some non-coding AD SNPs contribute to AD pathogenesis.

## Background

Alzheimer’s disease (AD) is a neurodegenerative disease characterized by cognitive impairment. In postmortem brains from AD patients, amyloid beta (Aβ) deposits on the surface of neurons and intracellular aggregations of hyperphosphorylated tau protein are observed. The heritability of AD is estimated to be between 58% and 79% [1]. The *APOE* ε4 allele is the genetic factor with the strongest influence identified to date on the risk of late-onset AD (LOAD). Genome-wide association studies (GWASs) have been performed to search for other genetic factors associated with AD [2-7]. Recently, the International Genomics of Alzheimer’s Project (IGAP) conducted a meta-analysis of LOAD in a cohort of 74,046 individuals and reported 21 loci including additional susceptibility loci [8]. However, how the single-nucleotide polymorphisms (SNPs) identified in these GWASs for AD contribute to AD pathogenesis is poorly understood because most of these SNPs are located in non-coding regions, such as introns and intergenic regions. These AD-associated SNPs (AD SNPs) could be tag SNPs of surrounding functional exonic variants [9]; however, a fine-mapping study of *BIN1*, *CLU*, *CR1*, and *PICALM*, which are the closest genes to several AD SNPs, showed no direct association with AD pathogenesis [10].

Recent studies have reported that disease-associated non-coding SNPs alter the functions of regulatory sequences, such as enhancers that typically regulate gene expression levels. For instance, Soldner *et al*. showed that a non-coding risk variant rs356168, which is associated with Parkinson’s disease (PD), is located in an enhancer region and upregulates the expression level of a PD-susceptibility gene SNCA [11]. It is reported that some SNPs in AD influenced gene expression levels as in PD [12,13]. In particular, Karch *et al*. searched functional AD SNPs from 21 loci that were found in the IGAP GWAS and revealed that the *ZCWPW1* and the *CELF1* loci were associated with some gene expressions [13].

The SNPs that influence gene expression levels as mentioned above are called expression quantitative trait loci (eQTLs). eQTLs are useful for considering function of non-coding SNPs, however this approach only achieves indirect evidence because eQTL effects are usually determined by correlations between genotypes and expression levels of target genes [14]. One of the molecular mechanisms to explain eQTL effects is contact between eQTLs and target genes by the formation of chromatin loops. Chromatin regions including eQTLs fold in order to bring in proximity to the genes they regulate. A growing body of evidence indicates that disease-associated variants in enhancers affect the expression levels of distal genes via chromatin loops in several diseases such as frontotemporal lobar degeneration, which belongs to the group of neurodegenerative diseases that includes AD [15-18]. These findings suggest that non-coding AD SNPs may alter the functions of regulatory sequences, such as enhancers that typically regulate gene expression levels via chromatin loops. Thus, we hypothesized that non-coding AD SNPs are located in enhancers and affect gene expression levels.

To test this hypothesis, we analyzed 392 AD SNPs located in non-coding regions by integrating enhancer activity data and chromatin interaction data. In particular, we used data from the Encyclopedia of DNA Elements (ENCODE) project [19] and the Roadmap Epigenomics project [20]. These projects measured epigenomic markers, including histone modifications and DNase I-hypersensitive sites, across every human tissue or cell type, and used these data to estimate genome-wide chromatin states (e.g., whether an enhancer is activated or not) [21,22]. To identify chromatin–chromatin interactions such as chromatin loops, we used data from the chromosome conformation capture (3C) variant Hi-C, which can capture genome-wide chromatin interactions via high-throughput sequencing. We found that nearly 30% of the non-coding AD SNPs were located in enhancers and that they affected the expression of genes associated with Aβ clearance, synaptic transmission, and immune responses. Among the non-coding AD SNPs, rs1476679 at the *ZCWPW1* gene locus and rs7364180 at the *CCDC134* gene locus were associated with several expression quantitative trait locus (eQTL) genes. Finally, analysis of chromatin higher-order structure revealed direct associations between rs1476679 and eQTL genes. Our findings would explain the regulatory mechanism of this AD SNP.

## Results

### Nearly 30% of non-coding AD SNPs are located in enhancers

**Figure 1** provides an overview of our study. First, we collected AD SNPs from the GWAS catalog database [23]. These AD SNPs have GWAS p-values of less than 1.00 × 10^−6^, which is used as a suggested threshold in GWAS. Among the 406 AD SNPs, 392 SNPs (96.6%) were in non-coding regions, whereas the rest were missense and synonymous mutations **(Figure 2A)**. Next, we checked whether these non-coding AD SNPs were located in enhancers, using publicly available enhancer data. Enhancer locations were predicted based on 11 histone modifications and DNase I-hypersensitive sites quantified in 127 human tissues or cell types, including ten brain tissues (see Methods). We counted non-coding AD SNPs located in the enhancers that were predicted in one or more tissues or cell types. Among the 392 non-coding AD SNPs, 106 (27.0%) were in enhancers **(Figure 2B)**. Of these 106 SNPs, 40 (10.2% of the 392 non-coding AD SNPs) were in enhancers identified in one or more brain tissues.

**Figure 1.**
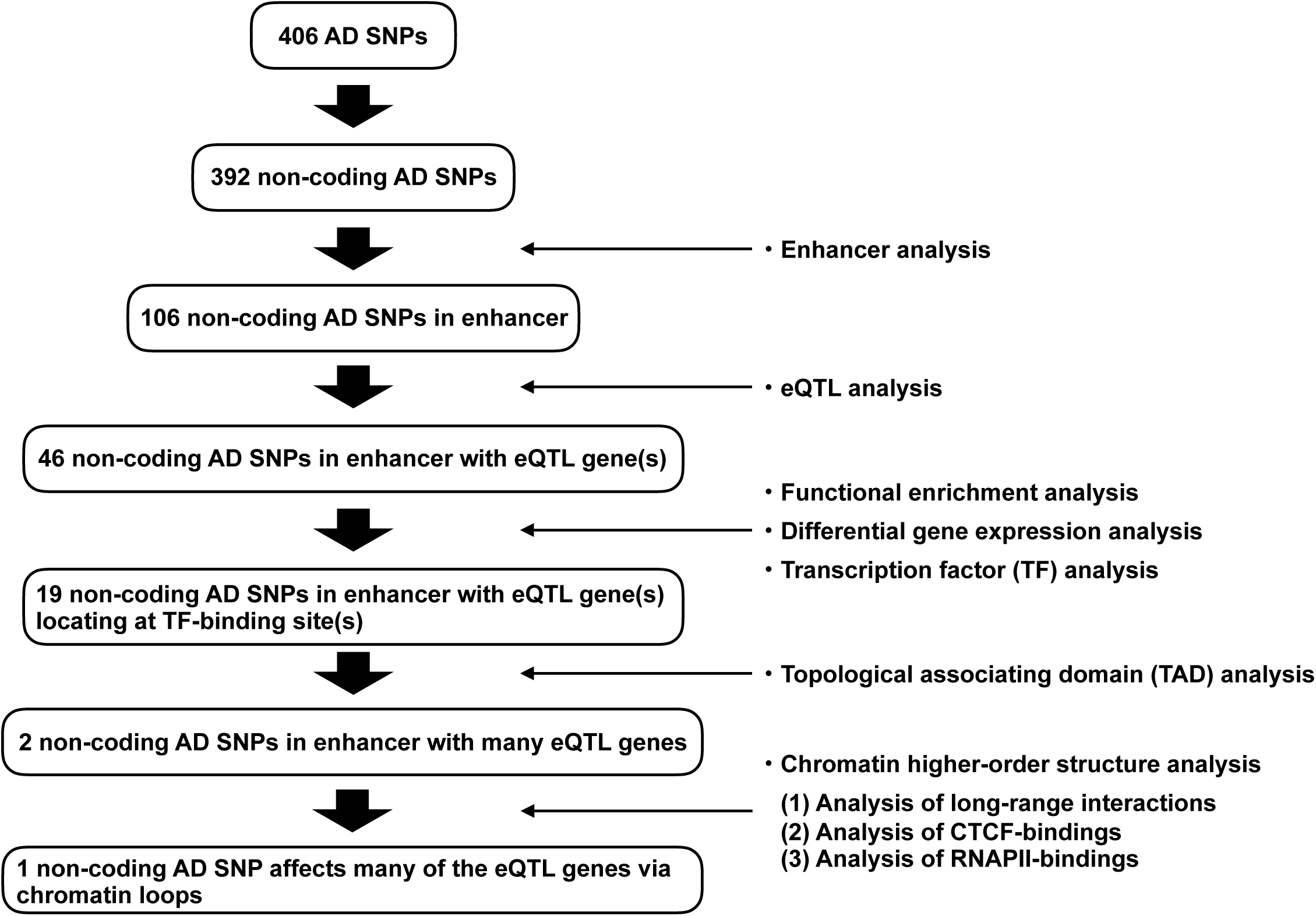
Flowchart of the present study.

**Figure 2.**
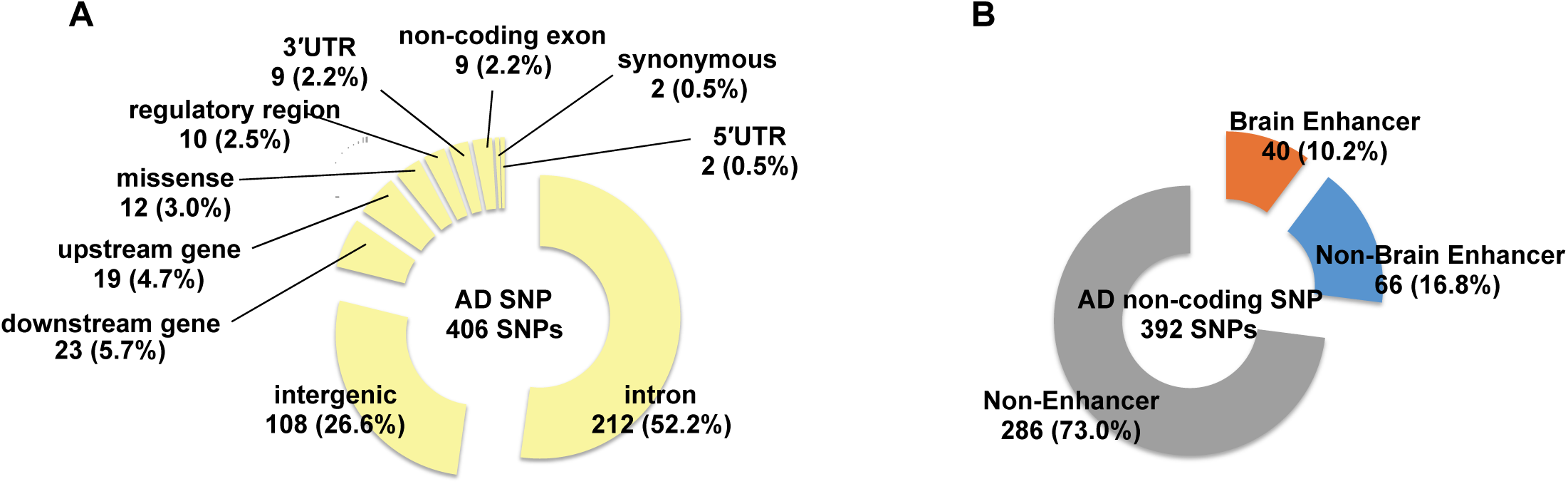
Nearly 30% of non-coding AD SNPs are located in enhancers. (A) Circle chart showing the genomic region location of AD SNPs from the GWAS catalog database (p-value < 1.00 × 10^−6^). (B) Circle chart showing the proportions of non-coding AD SNPs located in non-enhancer regions and in enhancers identified in one or more tissues or cell types. “Brain Enhancer” indicates non-coding AD SNPs located in enhancers identified in one or more brain tissues. “Non-Brain Enhancer” indicates non-coding AD SNPs located in enhancers that were not identified in brain tissues but were identified in the other tissues or cell types. All tissue and cell type names are described in Additional file 1: **Supplement Table S1**.

### Genes affected by non-coding AD SNPs are related to AD-relevant processes and are often differentially expressed in AD patients

Next, we investigated whether the non-coding AD SNPs functioned as eQTLs, which affect gene expression levels. To this end, the genes influenced by the non-coding AD SNPs (hereafter referred to as eQTL genes) were collected from the GTEx Portal [24,25] and BRAINEAC databases [26]. We used the eQTL genes that are located on the same chromosome as the associated AD SNPs. Among the 106 non-coding AD SNPs located in enhancers, 46 SNPs were associated with at least one eQTL gene and, overall, 130 eQTL genes were identified. These eQTL genes were related to Aβ formation, synaptic transmission, and immune responses **(Table 1)**. Interestingly, AD GWAS SNPs from a previous GWAS meta-analysis study were also associated with immune responses [27].

**Table 1.** Gene functional enrichment analysis.

| GO | Category | Description | Count | % | Log10(P) | Log10(q) |
| --- | --- | --- | --- | --- | --- | --- |
| GO:1902430 | GO Biological Processes | negative regulation of amyloid-beta formation | 3 | 2.38 | -5.03 | -0.94 |
| GO:0007271 | GO Biological Processes | synaptic transmission, cholinergic | 4 | 3.17 | -4.47 | -0.91 |
| R-HSA-1834949 | Reactome Gene Sets | Cytosolic sensors of pathogen-associated DNA | 5 | 3.97 | -4.34 | -0.89 |
| GO:0002768 | GO Biological Processes | immune response-regulating cell surface receptor signaling pathway | 11 | 8.73 | -4.29 | -0.89 |
| GO:0072665 | GO Biological Processes | protein localization to vacuole | 4 | 3.17 | -3.86 | -0.8 |
| GO:0006353 | GO Biological Processes | DNA-templated transcription, termination | 5 | 3.97 | -3.45 | -0.59 |
| GO:0016032 | GO Biological Processes | viral process | 13 | 10.32 | -3.35 | -0.53 |
| GO:0007169 | GO Biological Processes | transmembrane receptor protein tyrosine kinase signaling pathway | 11 | 8.73 | -2.74 | -0.26 |
| GO:0071466 | GO Biological Processes | cellular response to xenobiotic stimulus | 5 | 3.97 | -2.62 | -0.2 |
| GO:0007127 | GO Biological Processes | meiosis I | 4 | 3.17 | -2.36 | -0.02 |
| GO:0031329 | GO Biological Processes | regulation of cellular catabolic process | 11 | 8.73 | -2.33 | -0.02 |
| R-HSA-5653656 | Reactome Gene Sets | Vesicle-mediated transport | 10 | 7.94 | -2.31 | -0.01 |
| M254 | Canonical Pathways | PID MYC REPRESS PATHWAY | 3 | 2.38 | -2.27 | 0 |
| GO:0043547 | GO Biological Processes | positive regulation of GTPase activity | 7 | 5.56 | -2.09 | 0 |
| GO:0002274 | GO Biological Processes | myeloid leukocyte activation | 9 | 7.14 | -2.06 | 0 |

Terms with p-value < 0.01, minimum count 3, and enrichment factor > 1.5 (enrichment factor is the ratio between observed count and the count expected by chance) are collected and grouped into clusters based on their membership similarities.

We tested whether the eQTL genes were differentially expressed between AD and non-AD brain tissues. For this analysis, we used three publicly available datasets (syn5550404 [28], GSE5281 [29], and GSE44770 [30]) that include gene expression data analyzed in nine human brain tissues (prefrontal cortex, temporal cortex, visual cortex, entorhinal cortex, hippocampus, medial temporal gyrus, posterior cingulate, superior frontal gyrus, and cerebellum). Differentially expressed genes (DEGs) between AD and non-AD were identified in each brain tissue in each dataset (FDR<0.05) (Additional file 3: **Supplementary Table S3**)). We counted the number of the eQTL genes that were identified as the DEGs. To test whether the DEGs significantly include the eQTL genes, we performed 10,000 bootstrap replications in each brain tissue in each dataset and obtained the expected number of eQTL genes that are included in the DEGs. We calculated Z-scores and p-values from the number of eQTL genes and the expected numbers. Our results showed that the eQTL genes were significantly included among the DEGs in all these tissues with the exception of the cerebellum (Z-score > 0, p-value < 0.05) **(Table 2)**. The DEGs in the cerebellum had fewer the eQTL genes than them in the other brain tissues in the same dataset. These results are reasonable, as the cerebellum remains unaffected in Braak staging, which represents the pathological degree of AD [31]. Among the 126 eQTL genes analyzed in these datasets (4 of the 130 eQTL genes were not analyzed in the datasets because those corresponding probe sets were not constructed in the microarray or those transcripts did not satisfy criteria in RNA-seq), 110 genes (87.3%) were differentially expressed in one or more brain tissues or datasets. Additionally, 35 of 46 SNPs (76.1%) had one or more these differentially expressed eQTL genes. These results suggested that the non-coding AD SNPs affected genes whose expression levels were altered in the AD brain.

**Table 2.**
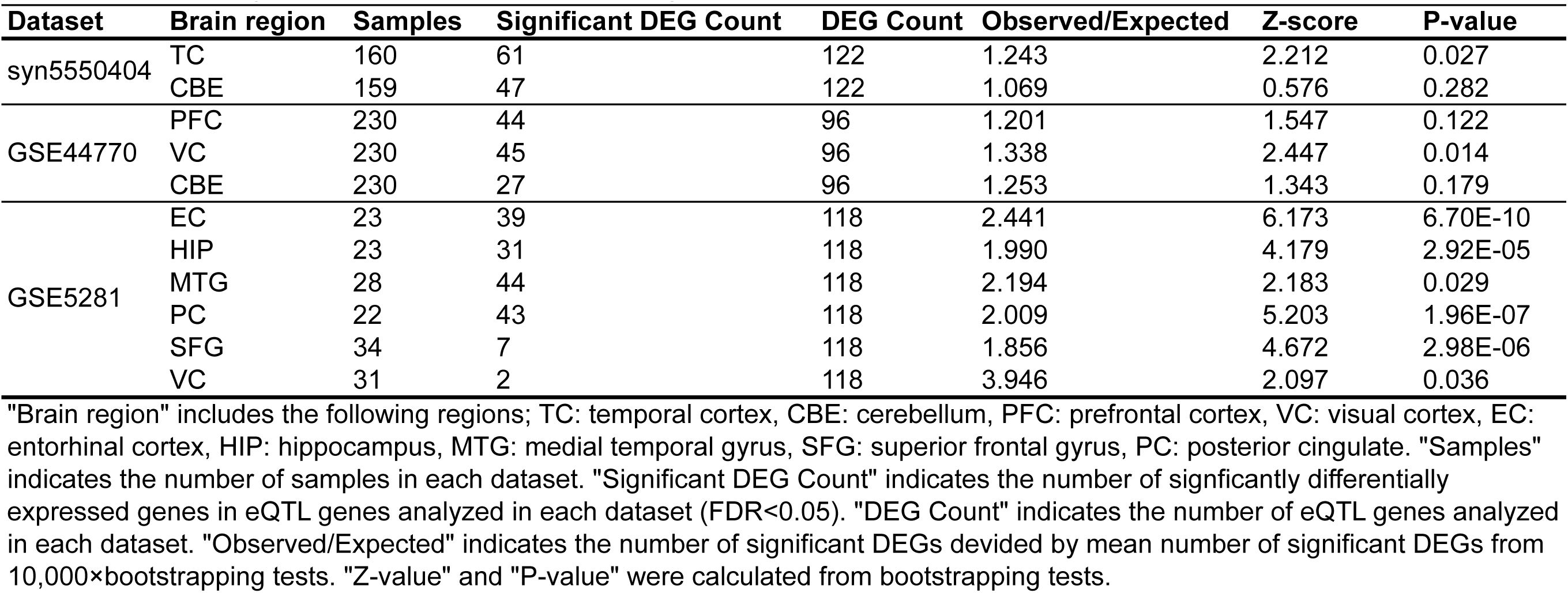
Bootstrapping test for DEG enrichment in eQTL genes.

### AD SNPs with eQTL effects are often located at protein-binding sites

Enhancers are regulatory regions that control the expression levels of surrounding genes when bound by specific proteins, such as transcription factors (TFs). To emphasize that the non-coding AD SNPs are located in the enhancers, we looked for TF-binding sites in these enhancers using the ENCODE ChIP-seq data for 161 TFs from 91 human cell types, which included 17 brain tissues or cell types (Additional file 4: **Supplementary Table S4**). Among the 46 SNPs with eQTL effects discussed above, 19 were located at a protein-binding site in at least one cell types **(Table 3)**. The closest genes were the corresponding eQTL genes for only eight of these SNPs, indicating that GWAS SNPs do not always affect the closest genes (**Table 3**). Four SNPs of the SNPs (rs4663105, rs1532278, rs4147929, and rs439401) were located around well-known AD candidate genes (*BIN1*, *CLU*, *ABCA7*, and *APOE*) and were eQTLs of those genes. The *BIN1* locus rs4663105 was located in enhancer that was activated in five tissues or cell types. Interestingly, all of these tissues or cell types were from brain regions including the hippocampus, suggesting that rs4663105 has the brain-specific eQTL effects (Additional file 5: **Supplement Table S5**). An enhancer near *CLU* locus was activated in 63 tissues or cell types including 4 brain tissues. The *APOE* locus rs439401 is located in the *APOE*-*APOC1* intergenic region. Enhancers near rs439401 were activated in 102 tissues or cell types including 7 brain tissues (Additional file 5: **Supplement Table S5**). This region is known as multienhancer 1 and affects *APOE* expression in various tissues or cell types, including macrophages, adipose tissue, and a neuronal cell line [32,33]. Indeed, *APOE* was identified as one eQTL gene of rs439401 in our study. On the other hand, 28 tissues or cell types where the enhancer involving the *ABCA7* locus was activated did not include brain tissues and were mainly from immune cells, such as monocytes, B cells, and T cells.

**Table 3.**
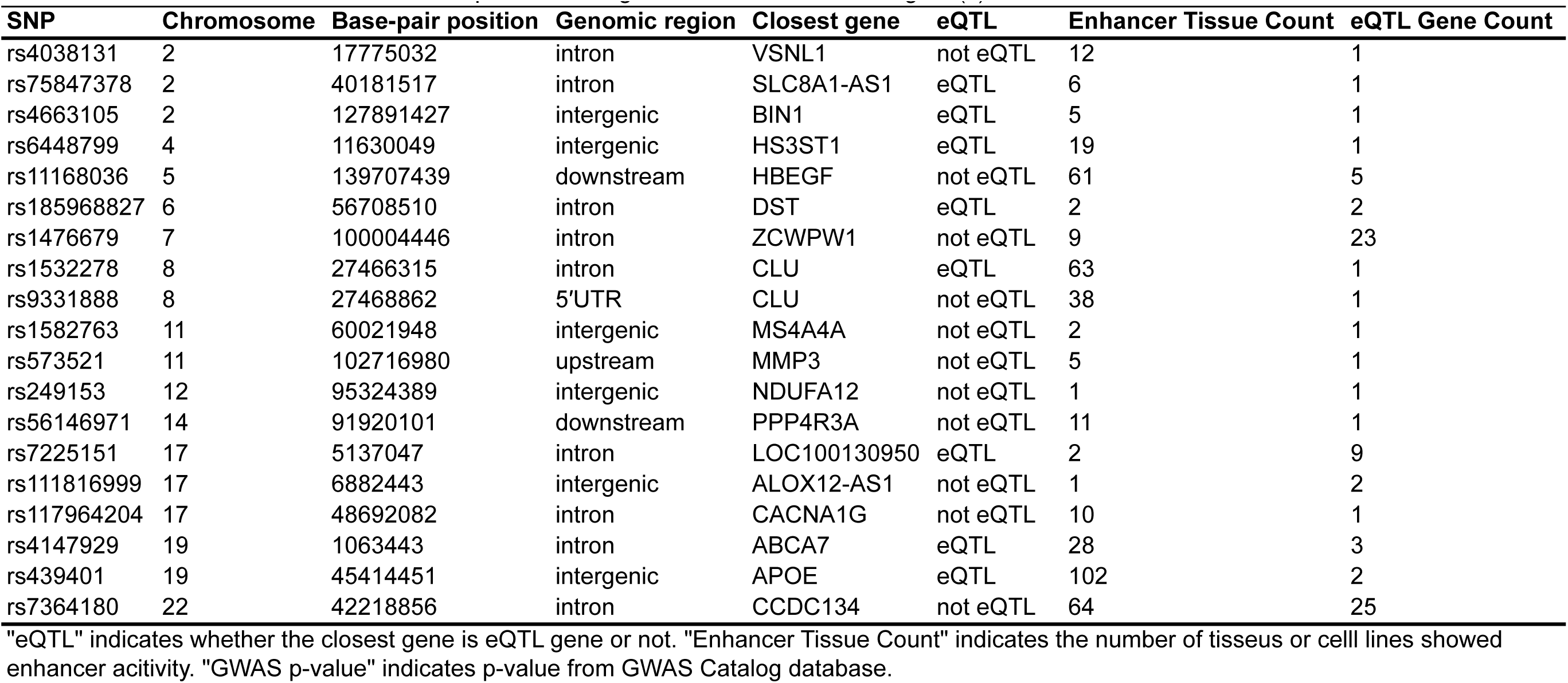
List of 19 SNPs that were located at protein-binding sites and that have eQTLgene(s).

| SNP | Chromosome | Base-pair position | Genomic region | Closest gene | eQTL | Enhancer Tissue Count | eQTL Gene Count |
| --- | --- | --- | --- | --- | --- | --- | --- |
| rs4038131 | 2 | 17775032 | intron | VSNL1 | not eQTL | 12 | 1 |
| rs75847378 | 2 | 40181517 | intron | SLC8A1-AS1 | eQTL | 6 | 1 |
| rs4663105 | 2 | 127891427 | intergenic | BIN1 | eQTL | 5 | 1 |
| rs6448799 | 4 | 11630049 | intergenic | HS3ST1 | eQTL | 19 | 1 |
| rs11168036 | 5 | 139707439 | downstream | HBEGF | not eQTL | 61 | 5 |
| rs185968827 | 6 | 56708510 | intron | DST | eQTL | 2 | 2 |
| rs1476679 | 7 | 100004446 | intron | ZCWPW1 | not eQTL | 9 | 23 |
| rs1532278 | 8 | 27466315 | intron | CLU | eQTL | 63 | 1 |
| rs9331888 | 8 | 27468862 | 5'UTR | CLU | not eQTL | 38 | 1 |
| rs1582763 | 11 | 60021948 | intergenic | MS4A4A | not eQTL | 2 | 1 |
| rs573521 | 11 | 102716980 | upstream | MMP3 | not eQTL | 5 | 1 |
| rs249153 | 12 | 95324389 | intergenic | NDUFA12 | not eQTL | 1 | 1 |
| rs56146971 | 14 | 91920101 | downstream | PPP4R3A | not eQTL | 11 | 1 |
| rs7225151 | 17 | 5137047 | intron | LOC100130950 | eQTL | 2 | 9 |
| rs111816999 | 17 | 6882443 | intergenic | ALOX12-AS1 | not eQTL | 1 | 2 |
| rs117964204 | 17 | 48692082 | intron | CACNA1G | not eQTL | 10 | 1 |
| rs4147929 | 19 | 1063443 | intron | ABCA7 | eQTL | 28 | 3 |
| rs439401 | 19 | 45414451 | intergenic | APOE | eQTL | 102 | 2 |
| rs7364180 | 22 | 42218856 | intron | CCDC134 | not eQTL | 64 | 25 |

"eQTL" indicates whether the closest gene is eQTL gene or not. "Enhancer Tissue Count" indicates the number of tissues or cell lines showed enhancer activity. "GWAS p-value" indicates p-value from GWAS Catalog database.

### AD SNPs and their eQTL genes co-localize in topologically associating domains

A fundamental mechanism underlying the effects of eQTLs on their regulated genes is enhancer-promoter regulation via chromatin higher-order structures, such as chromatin loops. Therefore, we examined whether the non-coding AD SNPs in enhancers regulate their eQTL genes through chromatin higher-order structures. We focused on topologically associated domains (TADs), which are genomic regions that spatially interact with each other (**Figure 3**) [34,35], since enhancers and their targeted promoters aggregate in the same TAD [36,37]. Therefore, we examined whether the 19 SNPs shown in **Table 3** co-localized with the corresponding eQTL genes in the same TAD. To detect TADs, we performed tethered conformation capture (TCC), which is a variant of the Hi-C method that is used for the identification of comprehensive chromatin loops through paired-end sequencing [38]. The neuroblastoma cell line SK-N-SH and the astrocytoma cell line U-251 MG were analyzed in the TCC experiment. These cell lines were used as models of brain cells. TADs were detected in each cell line using HiCdat and HiCseg software [39,40]. Among the 19 SNPs, 18 SNPs (94.7%) co-localized with at least one eQTL gene in the same TAD in the SK-N-SH and/or U-251 MG cell line (Additional file 6: **Supplement Table S6**). Furthermore, 13 SNPs in SK-N-SH and 14 SNPs in U-251 MG co-localized in the same TAD with more than 80% of the eQTL genes associated with that particular SNP. These results suggested that the AD SNPs might regulate eQTL genes in the same TAD through chromatin higher-order structures.

**Figure 3.**
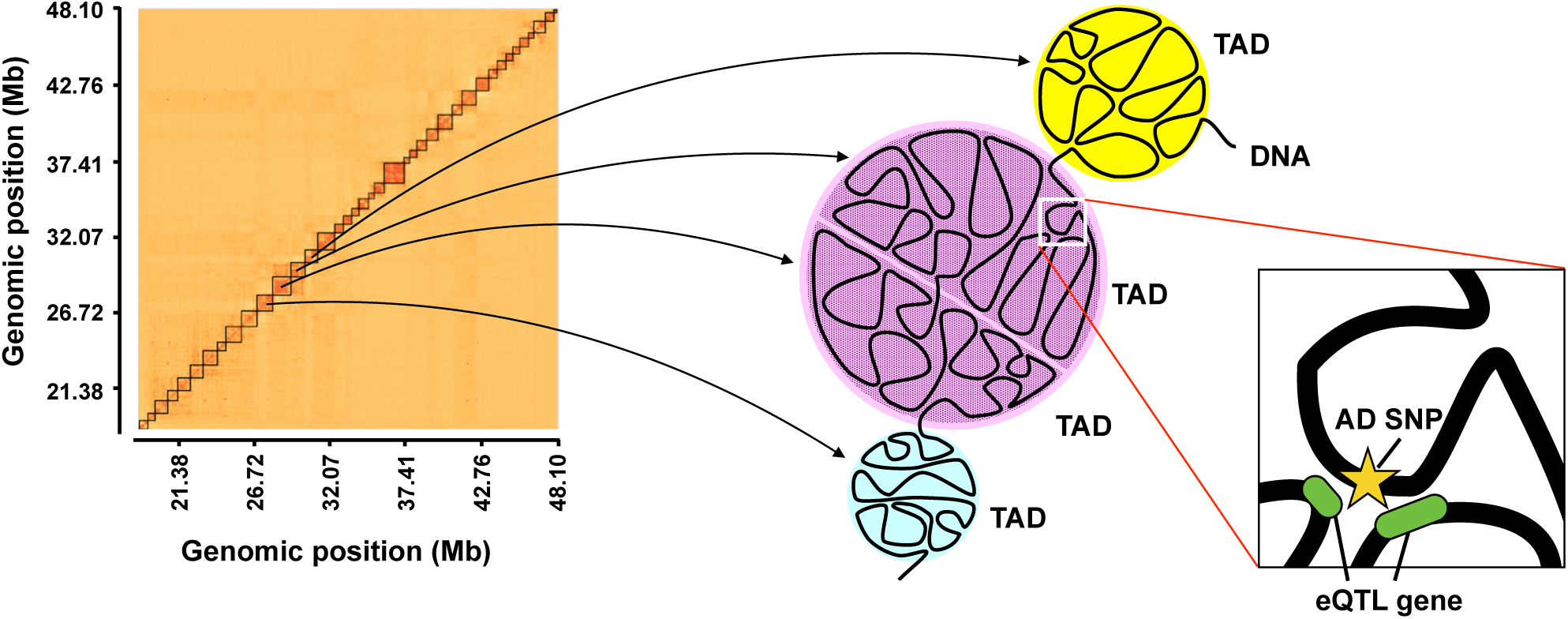
AD SNPs and their eQTL genes co-localize in topologically associating domains (TADs) Heatmap showing the frequency of chromatin interactions based on tethered conformation capture (TCC) experiments in the astrocytoma cell line U-251MG (100-kb bins). Diagonal darker blocks indicate TAD. AD SNP could contact the distal eQTL genes via chromatin interactions.

### AD SNPs associating with many eQTL genes are located in CTCF-binding sites

We found that rs1476679 and rs7364180 SNPs were associated with a particularly large number of eQTL genes (23 and 25, respectively) in **Table3**. rs1476679 is located in the intronic region of the *ZCWPW1* gene and the enhancer where it is located was activated in nine tissues and cell lines (**Figure 4A** and Additional file 7: **Supplementary Table S7**). rs7364180 is located in the intronic region of the *CCDC134* gene and in enhancer that was activated in 64 tissues or cell types (**Figure 4B** and Additional file 7: **Supplementary Table S7**). These SNPs co-localized with only approximately 30–70% (depending on the cell lines) of the corresponding eQTL genes in the same TAD (Additional file 6: **Supplement Table S6**), suggesting that these SNPs affected eQTL genes outside of the TADs via long-range chromatin interactions. Interestingly, rs1476679 and rs7364180 SNPs were localized at protein-binding sites of the CCCTC-binding factor (CTCF) in 12 and 66 cell lines, respectively, including neuronal cell lines (**Figure 4A,B** and Additional file 8: **Supplementary Table S8**). CTCF is a key factor to form chromatin loops and protects promoters against acting by chance from distant enhancers [41]. Chromatin loops are formed by the dimerization of two CTCF proteins binding to both regions that interact each other and the binding of a ring-shaped cohesin complex **(Figure 4C)** [42-44]. The formation of chromatin loops draws enhancers closer to promoters and can influence the expression of nearby or distant genes. In addition, we found binding sites for several TFs within the 1 kb region downstream from rs1476679 **(Figure 4A)** and in the region including rs7364180 **(Figure 4B)**. The TFs binding to these regions included SMC3 and RAD21, which are components of the cohesin complex. These findings suggested that rs1476679 and rs7364180 might be involved in the formation of chromatin loops via CTCF which could regulate the expression levels of eQTL genes in cooperation with nearby enhancer regions.

**Figure 4.**
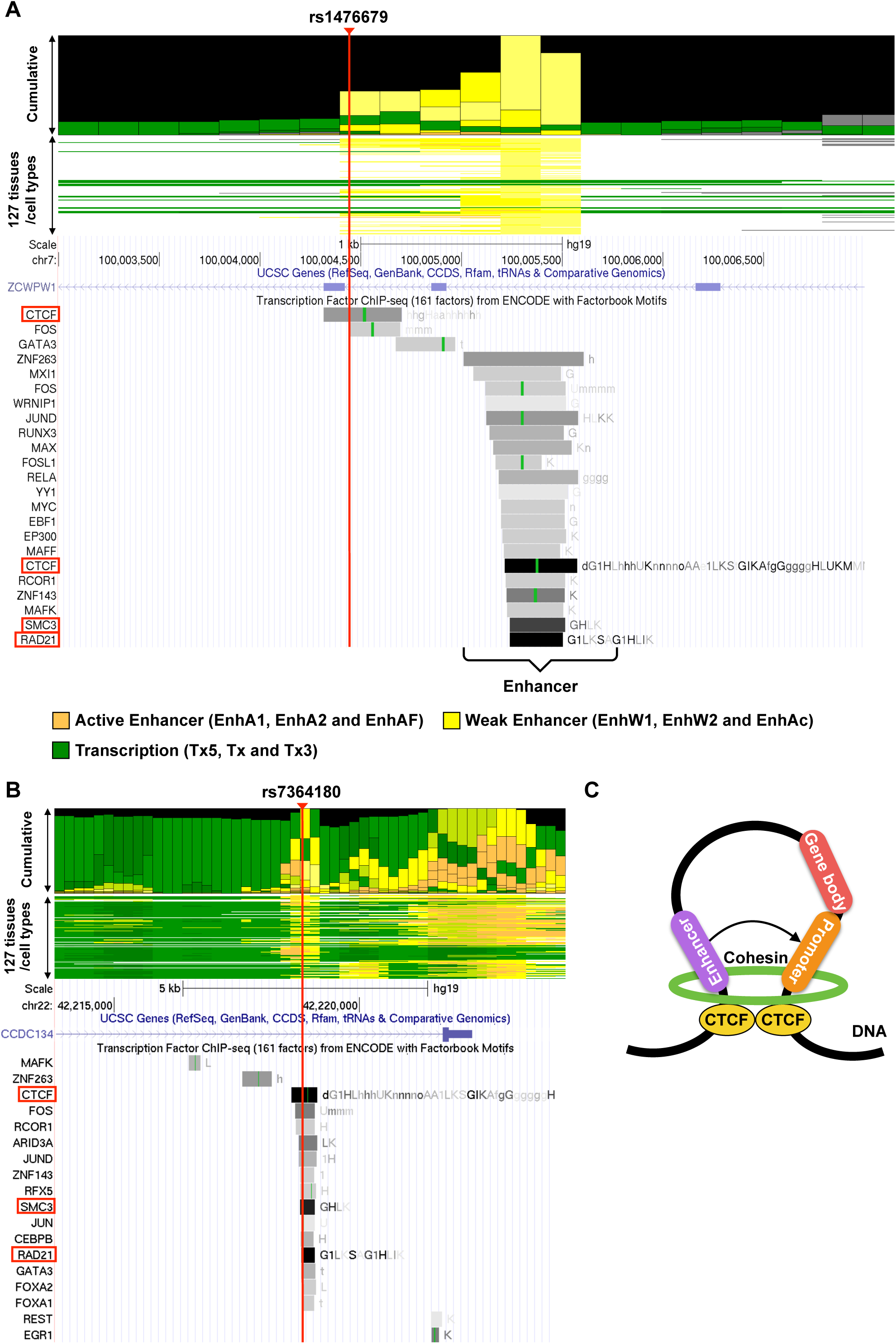
AD SNPs with eQTL effects are often located at protein-binding sites. (A, B) Cumulative bar graph of the chromatin state across 127 tissues or cell types (upper panel) and ChIP-seq tracks (bottom panel) around rs1476679 (A) and rs7364180 (B). Three representative chromatin state groups of the cumulative bar graph are depicted according to the color legend. The chromatin state names are shown in parentheses (see details in Methods). Details of all 25 chromatin state names are described in Additional file 2: **Supplement Table S2**. Grey bars in ChIP-seq tracks represent peak clusters of transcription factor (TF) occupancy. The color intensity of the bars is proportional to the level of TF occupancy. Green bars represent motif sites for the corresponding TFs. These ChIP-seq tracks were generated from the UCSC genome browser (https://genome.ucsc.edu/). (C) A schematic representation of a chromatin loop based on CTCF binding.

### rs1476679 spatially contacts many eQTL genes via CTCF-mediated chromatin loops

To provide further insight on how rs1476679 and rs7364180 may have an effect on eQTL genes through chromatin higher-order structures, we performed the following analyses: (1) investigation of whether rs1476679 and rs7364180 displayed long-range chromatin interactions; (2) evaluation of whether the SNPs and their eQTL genes spatially contacted each other through CTCF-CTCF interactions; and (3) examination of whether RNA polymerase II (RNAPII) bound upstream of the eQTL genes interacting with the SNPs.

First, we investigated chromatin loops formed by rs1476679. To this end, we applied the fourSig method [45] to the TCC data from the SK-N-SH and U-251 MG cell lines. We found that the chromatin loops extended approximately ±500 kb from rs1476679 **(Figure 5A)**. We analyzed publicly available data and validated this extensive interacting region through chromatin interaction analysis using paired-end tag sequencing (ChlA-PET), which is experimental method used to identify chromatin loops, in the 3D Genome Browser [46] (Additional file 9: **Supplement Figure S1**). The identified chromatin loops were located within 5 kb of the transcription start sites (TSSs) of 15 eQTL genes associated with rs1476679 (15 out of 23 genes = 65.2%), suggesting that rs1476679 spatially contacted many of the eQTL genes through long-range chromatin interactions.

**Figure 5.**
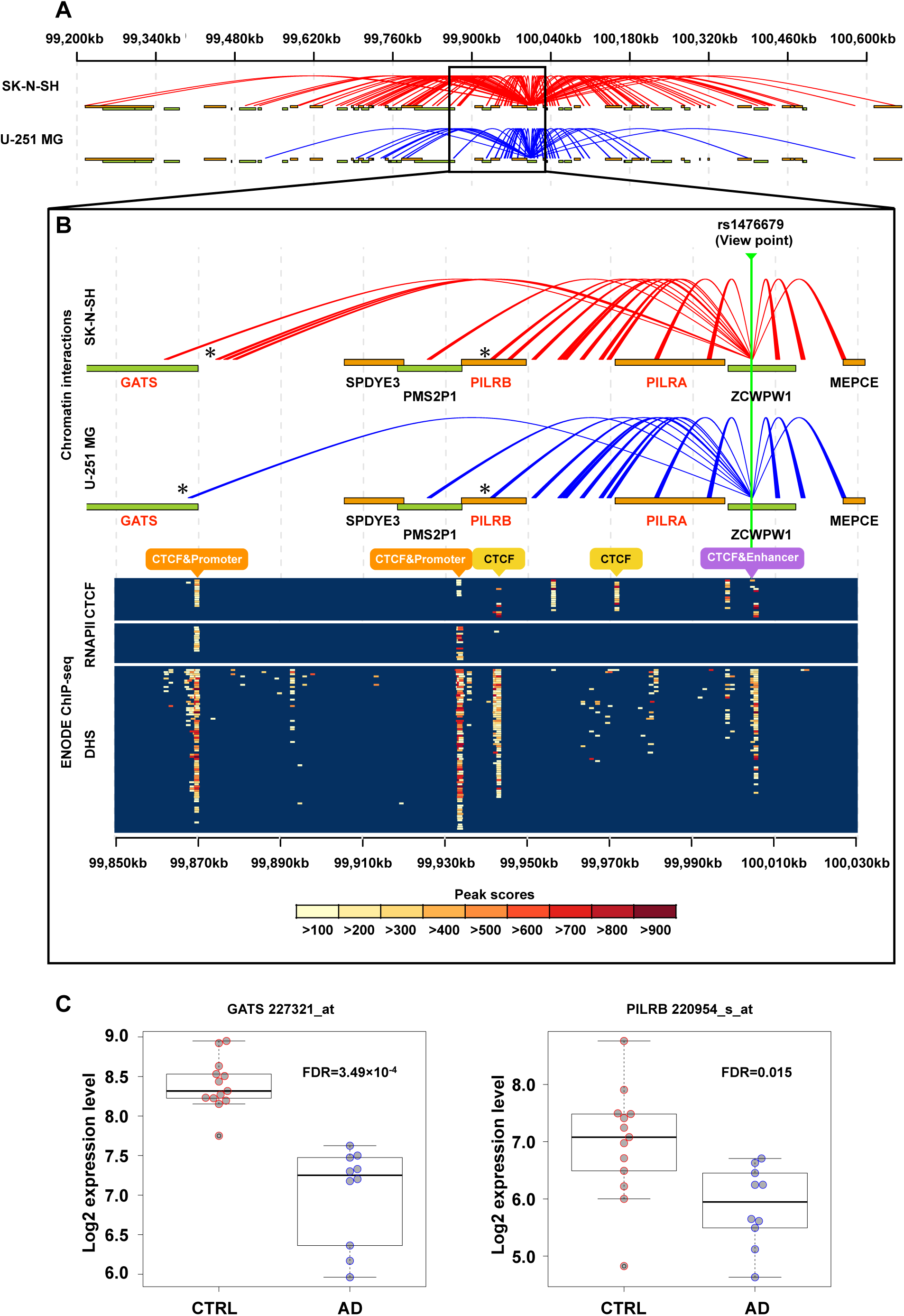
rs1476679 spatially contacts many eQTL genes via CTCF-mediated chromatin loops and affects their expression levels. (A) Chromatin interactions of the rs1476679 locus as determined by TCC experiments. Red and blue lines represent significant chromatin interactions from rs1476679 in SK-N-SH and U-251 MG cells, respectively. (B) Zoom-in region of the rs1476679 locus. The upper panel indicates chromatin interactions of the s1476679 locus. The orange and green bands indicate gene bodies on the positive and negative strands, respectively. Gene symbols in red indicate the eQTL genes of rs1476679. Asterisks indicate chromatin interactions of the rs1476679 locus with the *GATS*, *PILRB*, and *PILRA* genes. In the bottom panel, the color plot indicates the peak scores from ChIP-seq data for CTCF or RNA polymerase II (RNAPII) and DNase-seq data showing DNase I-hypersensitive sites (DHSs). Each row in the colored plot represents different brain tissues or neuronal cell lines (18 experiments (rows) including nine tissues or cell lines based on CTCF ChIP-seq, 21 experiments including ten cell lines based on RNAPII ChIP-seq, and 82 experiments including 31 cell lines based on DNase-seq; Additional file 10: **Supplement Table S9**). (C) *GATS* and *PILRB* expression levels in the hippocampus from GSE5281. Boxes represent the interquartile range between the first and third quartiles and median (internal line). Whiskers denote the lowest and highest values within 1.5 times the range of the first and third quartiles, respectively; dots represent *GATS* and *PILRB* expression levels in each sample.

Previous eQTL studies of AD have indicated that rs1476679 was associated with *PILRB* and *GATS* gene expression levels [13,15]. To examine whether rs1476679 contacts promoter regions of these genes via chromatin loops, we visualized significant chromatin loops with the eQTL genes *PILRB* and *GATS* genes, which were significantly downregulated in the AD hippocampus (FDR = 3.49 × 10^−4^ for *GATS* and FDR = 0.015 for *PILRB*) **(Figure 5B (upper panel) and Figure 5C)**. In particular, the rs1476679 region significantly interacted with the gene body of *PILRB* and the upstream region of *GATS* in the two cell lines analyzed (**asterisks in Figure 5B (upper panel)**). We also found chromatin loops with the eQTL gene *PILRA* gene.

Next, we investigated whether the chromatin loops between rs1476670 and *PILRB* and *GATS* occurred via CTCF-CTCF interactions, which require binding of CTCF to each interacting region. To this end, we determined whether CTCF binds to the *PILRB* and *GATS* loci using ChiP-seq data for CTCF in brain tissues or neuronal cell lines from the ChIP-Atlas database (nine tissues or cell lines; Additional file 10: **Supplement Table S9**). Furthermore, we used DNase-seq data to identify CTCF-binding at DNase I-hypersensitive sites (DHSs) (i.e., open-chromatin regions) (31 tissues or cell lines; Additional file 10: **Supplement Table S9**). The bottom panel in **Figure 5B** shows the peak scores for CTCF-binding sites and DHSs. As expected, we recognized CTCF-binding sites and DHSs in the rs1476679 region. Additionally, we found CTCF-binding sites in the gene body of *PILRB* and upstream of *PILRB* and *GATS*. These results suggested that rs1476679 spatially interacts with *PILRB* and *GATS* via CTCF.

Besides CTCF various other TFs were found to bind to the enhancers in the rs1476679 region and the region 1 kb downstream (**Figure 4A**), suggesting that these TFs could act on promoter regions of *PILRB* or *GATS* located in the rs1476679-interacting regions. To assess this hypothesis, we looked for RNAPII-binding promoter regions upstream of *PILRB* and *GATS*, in the rs1476679-interacting region, using ChIP-seq data for RNAPII in brain tissues or neuronal cell lines from the ChIP-Atlas database (ten tissues or cell lines, Additional file 10: **Supplement Table S9**). By combining the ChIP-seq data with the DNase-seq data mentioned above, we identified two regions upstream of *PILRB* and *GATS* that included both RNAPII-binding sites and DHSs, indicating that these two regions are active promoter regions in neuronal cell lines (**Figure 5B (bottom panel)**). The presence of active promoters in these regions was consistent with estimations based on histone modifications (Additional file 11: **Supplement Figure S2**). Taken together, our results suggested that the enhancers near rs1476679 approached the promoter regions of *PILRB* and *GATS* via CTCF-CTCF interactions.

We visualized significant chromatin loops with the region 100 kb downstream of rs1476679 to search for other candidate eQTL genes affected by chromatin loops from rs1476679. We found that rs1476679 interacted with a region within approximately 6 kb of the TSS of the *NYAP1* (neuronal tyrosine-phosphorylated phosphoinositide-3-kinase adaptor 1) gene (Additional file 12: **Supplement Figure S3B**), which showed a strong eQTL effect with rs1476679 (p-value = 1.16 × 10^−11^ in adipose subcutaneous tissues in the GTEx database), and was significantly upregulated in the AD hippocampus (FDR = 1.35 × 10^−4^) (Additional file 12: **Supplement Figure S3C**). CTCF binding sites and active promoter peaks were found in the region upstream of *NYAP1* (Additional file 12: **Supplement Figure S3B**, bottom panel). These results suggested that rs1476679 affects *NYAP1* expression via CTCF-CTCF interactions.

Taken together, our results from the chromatin higher-order structure analysis showed that rs1476679 spatially contacted several eQTL genes via chromatin loops and that rs1476679 likely affects *PILRB* and *GATS*, which were reported as the eQTL genes of rs1476679 in previous studies, through enhancer-promoter interactions. These enhancer-promoter interactions were supported by bindings of various TFs near the rs1476679 region and bindings of RNAPII in upstream of *PILRB* and *GATS*.

### The impact of rs7364180 on many of its eQTL genes may be indirect

Finally, we used a similar analysis to identify chromatin loops formed by rs7364180. We found that rs7364180 significantly interacted with *CCDC134* and its adjacent genes *MEI1* and *SREBF2* (Additional file 13: **Supplement Figure S4A**); however, no long-range chromatin interactions with the other eQTL genes were
identified. These results suggested that rs7364180 does not directly influence the expression levels of most of its eQTL genes. However, *SREBF2* showed strong eQTL effects with rs7364180 in several brain tissues (Additional file 13: **Supplement Figure S4B**). To examine the genes that are regulated by *SREBF2*, whose product is a TF, we used TRRUST, which is a TF-target interaction database based on text mining and manual curation [47]. This analysis showed that *SREBF2* regulates 20 genes that are significantly associated with AD (FDR = 1.60 × 10^−6^) (Additional file 14: **Supplement Table S10**, Additional file 15: **Supplement Table S11**). Therefore, many of the eQTL genes identified for rs7364180 may be indirectly affected by the change in *SREBF2* expression.

## Discussion

Previous GWASs have found AD-candidate SNPs, however, how these AD SNPs act to the pathogenesis is little known. In this study, we attempted to uncover those functions, considering epigenetic effects from chromatin higher-order structure. We confirmed our hypothesis that many non-coding AD SNPs are located in enhancers and affected the expression levels of surrounding genes. We also investigated chromatin higher-order structure with the aim of revealing direct interactions between the AD SNPs and eQTL genes through TCC experiments. We report the following findings: (1) nearly 30% of non-coding AD SNPs are located in enhancers; (2) the eQTL genes of the non-coding AD SNPs within enhancers are associated with Aβ formation, synaptic transmission, immune responses, and AD status; (3) rs1476679 and rs7364180 are associated with a particularly large number of eQTL genes; and (4) rs1476679 spatially interacts with many eQTL genes via chromatin loops.

Our findings revealed that rs1476679 is not only found in the enhancer but also directly interacts with eQTL genes through chromatin loops. In addition to *PILRB* and *GATS*, which were reported in previous studies [13,15], we found *NYAP1* to be a candidate eQTL gene affected by rs1476679 via a chromatin loop. *NYAP1* regulates neuronal morphogenesis [48]. A recent large-scale GWAS of AD identified an SNP around *NYAP1* [49] and we found *NYAP1* to be upregulated in the AD hippocampus. Thus, *NYAP1* may be related to AD pathology.

Our TCC experiments showed that rs7364180 interacts with *CCDC134* and its adjacent genes, *MEI1* and *SREBF2*. Although chromatin interactions with other
eQTL genes were not identified, we found that *SREFB2*, which is a TF, regulates the expression of 20 genes significantly associated with AD. In addition, previous studies have shown that *SREBF2* controls brain cholesterol synthesis and is involved in diabetes, which is associated with an increased risk of AD [50,51], and that AD model mice overexpressing *SREBF2* show Aβ accumulation and neurofibrillary tangle formation [52]. Overall, these findings suggest that rs7364180 might exert its effect on AD-associated genes, at least in part, indirectly via *SREBF2*.

rs1476679 and rs7364180 are located in CTCF-binding sites. CTCF is a regulator of chromatin topology that regulates the boundaries of TADs [53-56]. Mutations in CTCF-binding sites are associated with diseases [17,57]. For instance, in frontotemporal lobar degeneration, which belongs to the group of neurodegenerative diseases that includes AD, a SNP in a CTCF-binding site modifies the surrounding chromatin conformation and spatially regulates the expression level of a causative gene, *TMEM106B*, leading to neuronal death [57]. These reports support the hypothesis that the disease risk associated to rs1476679 and rs7364180 are due to epigenetic effects occurring via chromatin loops.

We found 19 SNPs in enhancer regions for which TF binding was confirmed by ChIP-seq data and that were associated with at least one eQTL gene **(Table 3)**. Of them, rs4147929 in the *ABCA7* intron was identified through IGAP GWAS [8]. The enhancer including the *ABCA7* locus was activated in immune cells, such as monocytes, B cells, and T cells. *ABCA7* is highly expressed in human monocytes that induced into macrophages [58]. Additionally, the expression level of *ABCA7* is also high in human microglia [59]. The monocytes-derived macrophages and microglia response to immune responses and have phagocytic activities. The epigenetic data that we used in this study did not include them from microglia, however, epigenetic status between the monocytes and microglia may similar. This suggests that the *ABCA7* locus rs4147929 could have the eQTL effects in microglia and could affect pathology in brain regions.

## Conclusions

In conclusion, multi-omics data analyses, including analyses of histone modifications, eQTL associations, protein binding, and chromatin higher-order structure data, suggested mechanisms by which non-coding AD SNPs identified in AD GWASs may confer disease risk. The main novel finding of this investigation is the eQTL mechanisms identified between rs1476679 at the *ZCWPW1* locus and its eQTL genes through chromatin interaction analysis. In future studies, we need to compare postmortem human brains from AD patients with those of normal healthy individuals to clarify the details of chromatin higher-order structure in AD.

## Methods

### AD-associated SNPs (AD SNPs)

AD SNPs were obtained from the GWAS catalog database (Release 20170627, https://www.ebi.ac.uk/gwas/) [23]. These SNPs included “Alzheimer” in the “DISEASE/TRAIT” column of the GWAS catalog data. We investigated 406 of these SNPs, including 19 confirmed SNPs identified in the IGAP study [8] and AD SNPs with GWAS p-values of less than 1.00 × 10^−6^, which is used as a suggested threshold in GWAS. The genomic positions of all SNPs were standardized to the human reference genome (hg19) based on their reference SNP ID (rsID). SNPs without a rsID were manually curated.

### Enhancer data from 127 tissues or cell types

A chromatin state model for 127 tissues or cell types was obtained from the Roadmap Epigenomics website (http://egg2.wustl.edu/roadmap/web_portal/). These 127 tissues or cell types are described in Additional file 1: **Supplementary Table S1**. The chromatin state model segments the human genome into 25 states based on 12 chromatin marks (H3K4me1, H3K4me2, H3K4me3, H3K9ac, H3K27ac, H4K20me1, H3K79me2, H3K36me3, H3K9me3, H3K27me3, H2A.Z, and DNase I-hypersensitive sites) using ChromHMM and ChromImpute [21,22]. We extracted six enhancer states (Active Enhancer 1 (EnhA1), Active Enhancer 2 (EnhA2), Active Enhancer Flank (EnhAF), Weak Enhancer 1 (EnhW1), Weak Enhancer 2 (EnhW2), and Primary H3K27ac possible Enhancer (EnhAc)) from the 25 states and treated them as enhancer data (Additional file 2: **Supplementary Table S2**). EnhA1, EnhA2, and EnhAF show high levels of H3K4me1 and H3K27ac, which are enhancer-associated histone modifications. EnhW1 and EnhW2 show high H3K4me1 and low H3K27ac levels. EnhAc shows low H3K4me1 and high H3K27ac levels.

### Expression quantitative trait loci (eQTLs)

The eQTL genes of each AD SNP were searched in the GTEx Portal database (https://www.gtexportal.org/) [24,25] and the BRAINEAC database (http://www.braineac.org/) [26]. For further details, see Additional file 16: **Supplementary Information**. The eQTL genes in the above databases are located on the same chromosome as the associated SNPs. Pseudogenes were removed based on the GENCODE pseudogene resource from the eQTL analysis [60]. The AD SNPs were considered to associate with the eQTL genes if the corrected p-value was less than 0.05. Each p-value was corrected for multiple testing across genes on the same chromosome using Storey’s q-value [61]. Gene functional enrichment analysis of the eQTL genes was performed using the Metascape database (http://metascape.org/) [62].

### Differentially expressed genes (DEGs) from publicly available datasets

DEGs between AD and non-demented brains were identified using three publicly available gene expression datasets (syn5550404 [28], GSE5281 **[29],** and GSE44770 [30]). For further details, see Additional file 16: **Supplementary Information**. The syn5550404 dataset contains RNA-seq data for cerebellum and temporal cortex samples from 312 Caucasian subjects with neuropathological diagnosis of AD, progressive supranuclear palsy, pathologic aging or elderly controls without neurodegenerative diseases. The DEGs were identified using multivariate linear regression after adjusting for covariates (age at death, gender, RNA integrity number (RIN), source, and flow cell). These statistics were provided by the AMP-AD Knowledge Portal (https://www.synapse.org/#!Synapse:syn2580853/wiki/409840). The GSE5281 dataset contains microarray data for six brain regions that are either histopathologically or metabolically relevant to 33 AD and 14 aging; these include the entorhinal cortex (BA 28 and 34), superior frontal gyrus (BA 10 and 11 and approximate BA 8), hippocampus, primary visual cortex (BA 17), middle temporal gyrus (BA 21 and 37 and approximate BA 22), and the posterior cingulate cortex (BA 23 and 31). The GSE44770 dataset contains microarray data for 230 autopsied tissues from dorsolateral prefrontal cortex (PFC), visual cortex (VC) and cerebellum (CR) in brains of LOAD patients, and non-demented healthy controls. These two datasets were reanalyzed, because statistics for the comparisons were not provided. The reanalyses of GSE5281 and GSE44770 was performed using t-tests and logistic regression analyses with covariates (age, gender, postmortem interval in hours, sample pH, RIN, sample
processing, and batch), respectively, as described in the original analyses. DEGs were defined based on an FDR-adjusted p-value < 0.05.

### Cell culture

We employed two cell lines for this study: the neuroblastoma cell line SK-N-SH (American Type Culture Collection, Manassas, VA, USA) and the astrocytoma cell line U-251 MG (Japan Collection of Research Bioresources Cell Bank, Ibaraki, Osaka, Japan). Both cell lines were grown in Eagle’s MEM and cultured at 37°C with 5% CO_2_. For further details, see Additional file 16: **Supplementary Information**.

### TCC library preparation and deep sequencing for target regions

Tethered conformation capture (TCC), which is a variation of Hi-C, was performed to detect chromatin interactions. A TCC library was prepared in accordance with the method reported by Kalhor *et al*. with minor modifications [38]. The captured DNA fragments corresponding to the target regions were obtained from the TCC library using the SureSelect Target Enrichment System (Agilent Technologies). The library was subjected to paired-end sequencing on the Genome Analyzer IIx or MiSeq (Illumina) platform. For further details, see Additional file 16: **Supplementary Information**.

### Processing the sequencing output

In accordance with the procedure established by Imakaev *et al*. [63], we mapped the sequenced reads to the human reference genome (hg19) using Bowtie2 and used the “hiclib” library (provided by the Leonid Mirny Laboratory (https://bitbucket.org/mirnylab/)) to filter out non-informative reads. For further details, see Additional file 16: **Supplementary Information**.

### Chromatin interaction analysis

To identify significant chromatin interactions, we applied the R software package fourSig [45]. We counted mapped reads from TCC to the nearest restriction sites (HindIII sites) because TCC assumes that chromatin interactions occur around restriction sites. A viewpoint nearest to the HindIII sites on both sides of the SNP was
selected when we detected chromatin interactions for an SNP. A window size of one fragment was set. The significance level employed was an FDR-adjusted p-value of 0.05.

### Identification of topologically associated domains (TADs)

Identification of TADs in the SK-N-SH and U-251MG cell lines was performed using the R software packages HiCdat and HiCseg [39,40]. The sequenced reads mapped to the human reference genome (hg19) were normalized using HiCdat and compiled using a genomic bin size of 100 kb. The default values for the HiCseg parameters were employed to detect TADs. HiCseg estimated the TAD block boundaries based on a maximum likelihood approach.

## Abbreviations

GWAS: Genome-wide association study
SNPs: Single-nucleotide polymorphisms
AD: Alzheimer’s disease
AD SNP: AD-associated SNP
eQTL: Expression quantitative trait locus
LOAD: Late-onset Alzheimer’s disease
DFG: Differentially expressed gene
TF: transcription factor
CTCF: CCCTC-binding factor
TCC: Tethered conformation capture
DHS: DNase I-hypersensitive site
RNAPII: RNA polymerase II
TAD: Topologically associated domain.

## Ethics approval and consent to participate

Not applicable.

## Consent for publication

Not applicable.

## Availability of data and materials

The datasets used and/or analysed during the current study are available from the corresponding author on reasonable request.

## Authors’ contributions

MK contributed to the study design. MK, NH and MH wrote the manuscript. MK analyzed the data. NH performed the high-throughput sequencing analysis. MH performed the chromatin higher-order experiments. MK, NH, MH, AM, RK, TI and AN contributed text to the manuscript. All authors read and approved the final manuscript.

## Acknowledgements

We acknowledge all groups that have contributed.

## Figure legends

**Figure S1.**
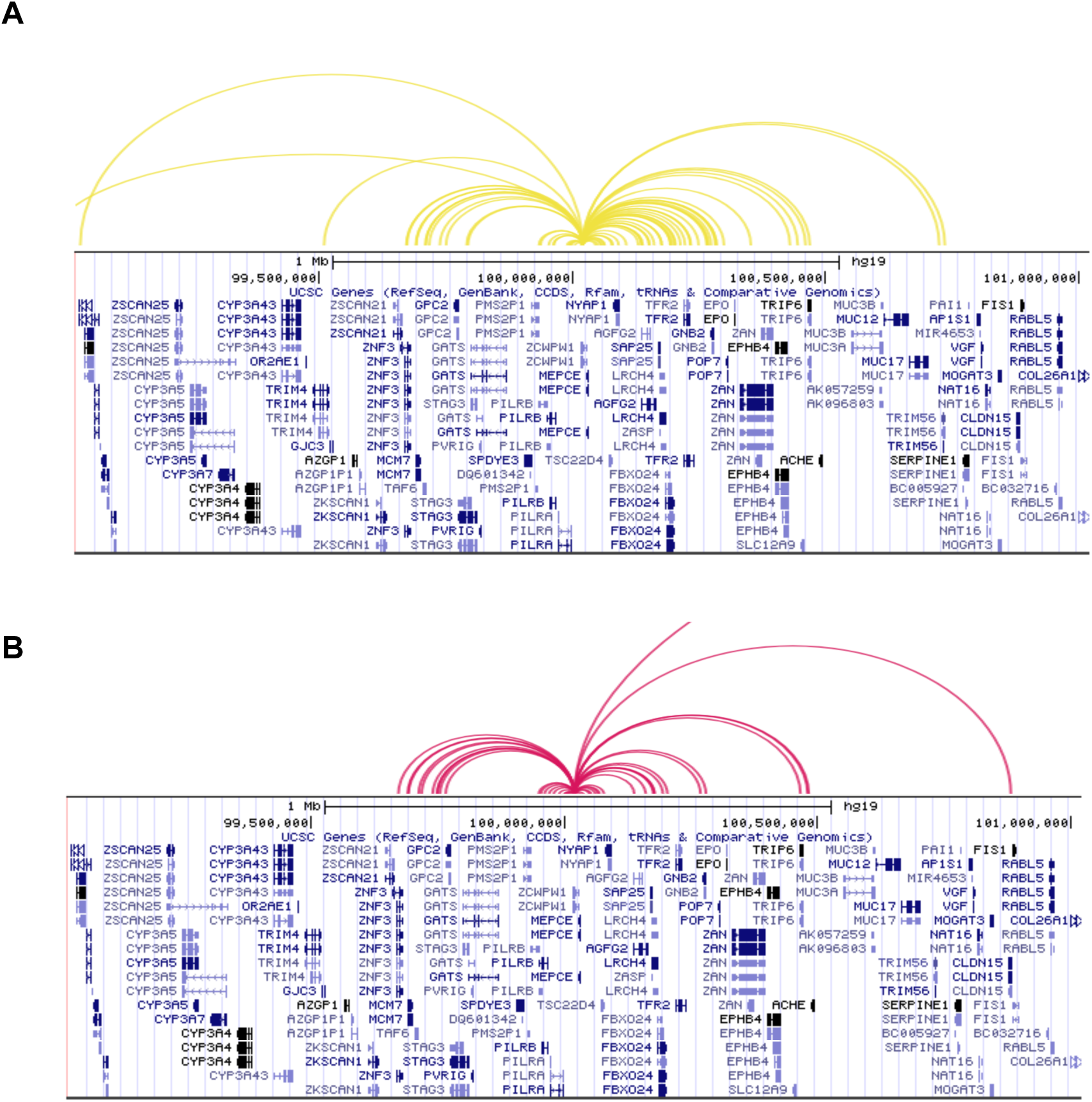
Higher-order chromatin structure of the rs1476679-containing region as assessed by chromatin interaction analysis by paired-end tag sequencing (ChIA-PET) experiments. Each curved line represents chromatin loops associated with RNA polymerase II in K562 (A) and MCF-7 (B) cell lines. These figures were modified from the 3D Genome Browser (http://promoter.bx.psu.edu/hi-c/index.html).

**Figure S2.**
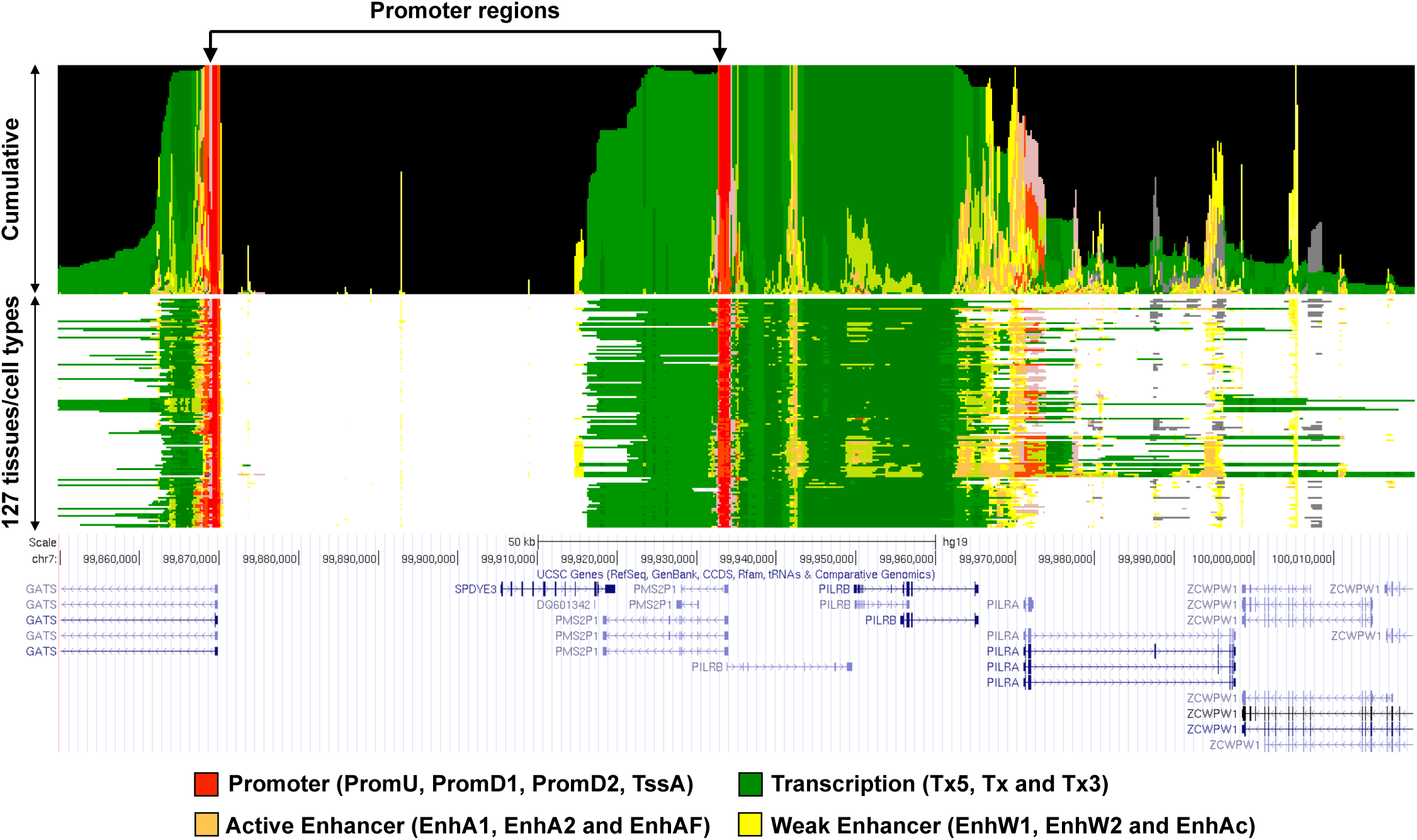
Upstream regions of *GATS* and *PILRB* genes show prominent promoter activity as estimated from histone modifications. Four representative chromatin state groups are shown (see color key). The chromatin state names are shown in parentheses (see detail in Materials and Methods). Details of all 25 chromatin state names are given in Additional file 2: **Supplement Table S2**.

**Figure S3.**
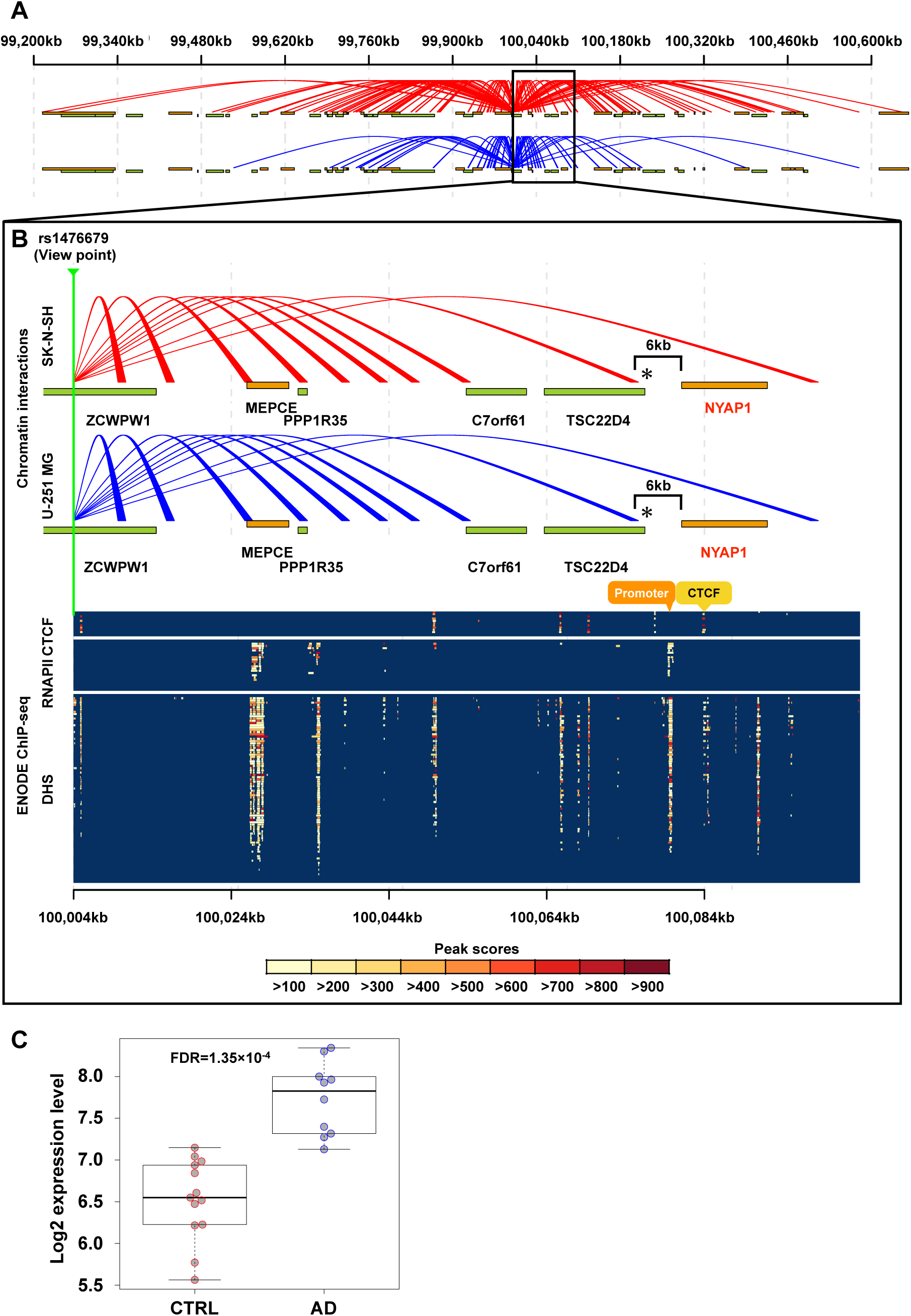
Chromatin interactions between the rs1476679 locus and *NYAP1*. (A) Chromatin interactions from rs1476679 locus as determined by TCC experiments. Red and blue lines represent statistically significant chromatin interactions from rs1476679 in SK-N-SH and U-251 MG, respectively. (B) Zoom-in region of rs1476679 locus. Upper panel indicates chromatin interactions from s1476679 locus. Orange and green bands indicate gene bodies on the positive and negative strands, respectively. Gene symbols in red indicate eQTL genes of rs1476679. Asterisks indicate chromatin interactions from the rs1476679 locus to around the TSS of *NYAP1*. In the bottom panel, a color plot indicates peak scores from ChIP-seq data for CTCF or RNA polymerase II (RNAPII) and DNase-seq data to show DNase I hypersensitive sites (DHSs). Each row in the color plot represents different neuronal cell lines (18 experiments (rows) including nine tissues or cell lines in CTCF ChIP-seq; 21 experiments including ten cell lines in RNAPII ChIP-seq; 82 experiments including 31 cell lines in DNase-seq; Additional file 10: **Supplement Table S9**). (C) Expression level of *NYAP1* in hippocampus from GSE5281. Boxes represent the interquartile range between the first and third quartiles and median (internal line). Whiskers denote the lowest and highest values within 1.5 times the range of the first and third quartiles, respectively; dots represent *GATS* and *PILRB* expression levels in each sample.

**Figure S4.**
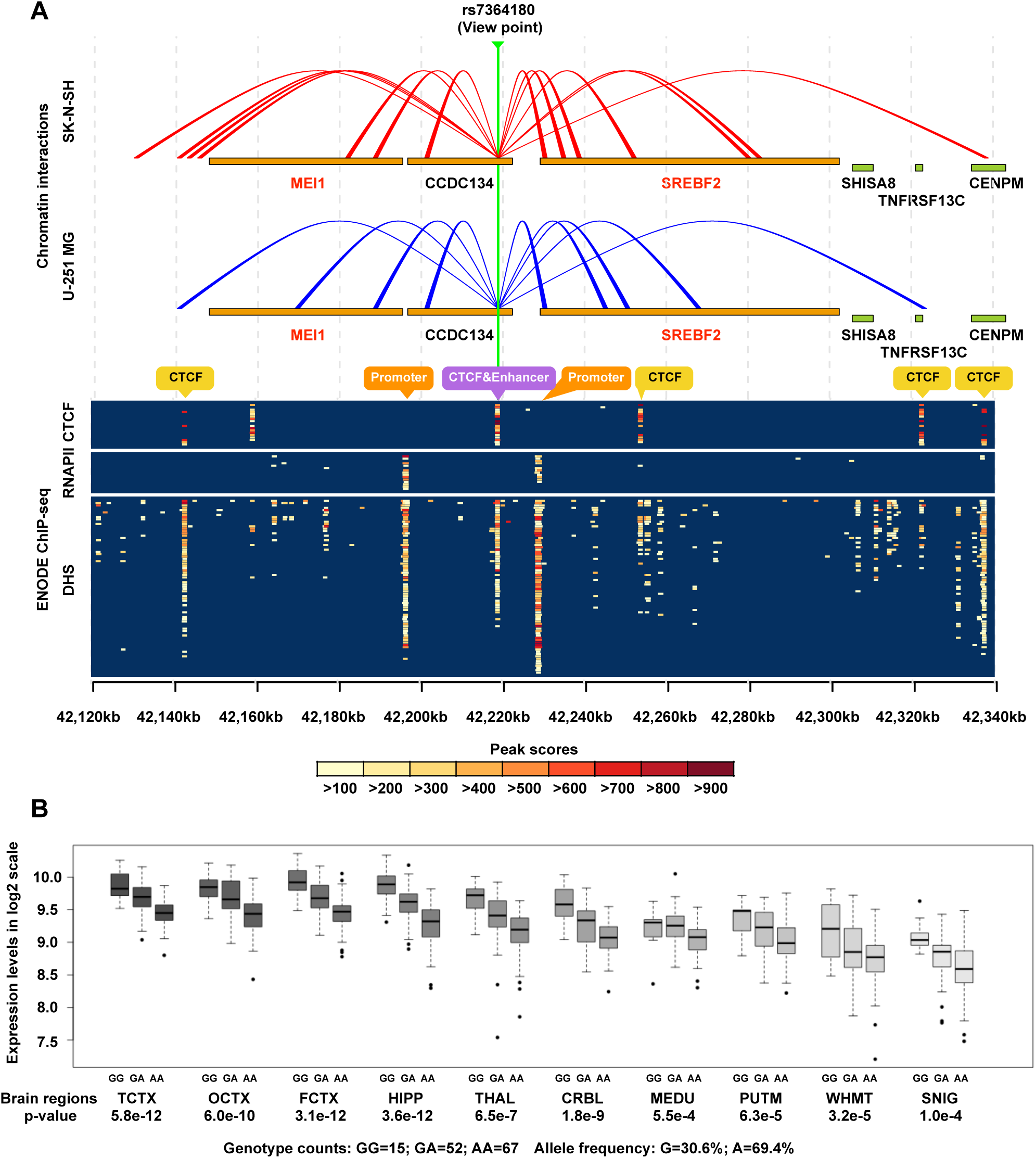
Higher-order chromatin structure of the rs7364180-containing region. (A) Chromatin interactions from the rs7364180 locus as determined by TCC experiments and protein binding in the corresponding region. In the upper panel, red and blue lines represent statistically significant chromatin interactions from rs7364180 in SK-N-SH and U-251 MG, respectively. Orange and green bands indicate gene bodies on the positive and negative strands, respectively. Gene symbols in red indicate eQTL genes of rs7364180. In the bottom panel, a color plot indicates peak scores from ChIP-seq data for CTCF or RNA polymerase II (RNAPII) and DNase-seq data to show DNase I hypersensitive site (DHS). Each row in the color plot represents different neuronal cell lines (18 experiments (rows) including nine tissues or cell lines in CTCF ChIP-seq; 21 experiments including ten cell lines in RNAPII ChIP-seq; 82 experiments including 31 cell lines in DNase-seq; Additional file 10: **Supplement Table S9**). (B) eQTL associations between rs7364180 genotypes (GG, GA, and AA) and *SREBF2* expression levels in the following ten brain tissues: TCTX, temporal cortex; OCTX, occipital cortex (specifically the primary visual cortex); FCTX, frontal cortex; HIPP, hippocampus; THAL, thalamus; CRBL, cerebellar cortex; MEDU, medulla (specifically the inferior olivary nucleus); PUTM, putamen; WHMT, intralobular white matter; and SNIG, substantia nigra. These box plots were modified from the BRAINEAC database.

## Supplementary Information

### Cell culture

We used two cell lines for this study; a neuroblastoma cell line, SK-N-SH (American Type Culture Collection, Manassas, VA, USA), and an astrocytoma cell line, U-251MG (Japan Collection of Research Bioresources Cell Bank, Ibaraki, Osaka, Japan). SK-N-SH was grown in Eagle’s MEM medium supplemented with 10% FBS, 1 mM sodium pyruvate, 0.1 mM each non-essential amino acids and antibiotics (Gibco(R) Penicillin-Streptomycin, 10,000 U/mL (Life Technologies)). U-251MG was grown in Eagle’s MEM medium supplemented with 10% FBS and antibiotics. Both cell lines were cultured at 37°C with 5% CO_2_.

### TCC library preparation

A TCC library was prepared according to a method reported by Kalhor *et al*. with minor modifications [1]. The cells were cross-linked with 2 % formaldehyde on 10 minutes incubation, and then 2 M glycine was added to a final concentration of 125 mM to stop the reaction. Since cells stuck strongly to the culture dish during the crosslinking reaction, which made it hard to harvest them, they were washed twice using PBS(-) and then treated with trypsin-EDTA (0.25% trypsin and 1 mM EDTA, Invitrogen) for 15 minutes at 37 °C. The dishes were chilled on ice and the cells were harvested by pipetting on ice to avoid proteolysis. To isolate nuclei, cells were suspended in 50 mL of ice-cold lysis buffer (10 mM Tris-HCl, pH 8.0, 10 mM NaCl, 1 % IGEPAL CA-630 (Sigma), and 1/500 (vol/vol) protease inhibitor cocktail (Nacalai Tesque)), stirred for 90 min at 4 °C and then centrifuged at 1,600× g [2,3]. Nuclei were suspended in wash buffer 1 (50 mM Tris-HCl, pH 8.0, 50 mM NaCl and 1 mM EDTA). The suspension was incubated for 10 minutes at 65 °C after the addition of 95 μL 2 % SDS. After that, 105 μL 25 mM iodoacetyl-PEG2-biotin (IPB) (Thermo Fisher Science) was added, followed by incubation for 40 minutes at room temperature to biotinylate cysteine residues. Chromatin was digested with *Hin*dIII (New England Biolabs) overnight at 37 °C with rotation. After digestion, the mixture was placed in a Slide-A-Lyzer Dialysis Cassette (20K MWCO) (Pierce Protein Research Products, Rockford, Illinois) and dialyzed for four hours at room temperature against 1 L of the TE buffer (10 mM Tris-HCl, pH 8.0, 1 mM EDTA) to remove IPB remaining from the biotinylation step. The TE buffer was renewed after three hours.

Digested chromatin was immobilized on 0.4 mL Dynabeads(R) MyOne Streptavidin T1 beads (Invitrogen) to remove non-crosslinked DNA fragments. The magnetic beads were treated with neutralized IPB and then washed twice with wash buffer 2 (10 mM Tris-HCl, pH 8.0, 50 mM NaCl, 0.4 % Triton X-100). The DNA ends were labeled by adding 0.7 μL 10 mM dATP, 0.7 μL 10 mM dTTP, 0.7 μL 10 mM dGTPalphaS (AXXORA), 15 μL 0.4 mM biotin-14-dCTP (Invitrogen), and 5 μL 5 units/μL Klenow (New England Biolabs), and then incubated at room temperature for 40 min. The labeling reaction was stopped by adding 5 μL 0.5 M EDTA, and then the beads were washed twice and resuspended in 500 μL of wash buffer 3 (50 mM Tris-HCl, pH 7.4, 0.4 % Triton X-100, 0.1 mM EDTA). For ligation, the beads suspension was added to 4.4 mL of ligation mix (containing 180 μL 10 % Triton X-100, 250 μL 10× ligase buffer (New England Biolabs), 100 μL 1 M Tris-HCl, pH 7.4, and 50 μL 100× BSA (New England Biolabs)). After the addition of 2 μL DNA ligase (New England Biolabs), the suspension was gently rocked on a reciprocal shaker at 16 °C. After 4 hours incubation, 0.2 mL 0.5 M EDTA was added to stop the ligation step. To purify the ligated DNA, reverse-crosslinking and protein digestion with proteinase K (New England Biolabs) were performed overnight at 65 °C. DNA free from streptavidin-coated beads was purified by phenol-chloroform-isoamylalcohol extraction. To determine whether or not the ligatin step had been successful, PCR amplification was performed using the primers described by Lieberman-Aiden *et al*. [4], and then the PCR products were sequenced using a Sanger sequencer ABI 3130 (Applied Bioscience).

After quality checking, biotinylated residues from unligated DNA ends were removed using *Escherichia coli* exonuclease III (New England Biolabs). The DNA was sheared with an acoustic solubilizer Covaris S2 (Covaris Inc.), and the fragment size was checked with a 2100 Bioanalyzer (Agilent Technologies). The biotin-labeled DNA fragments were pulled-down using 10 μL streptavidin-coated magnetic beads (Dynabeads(R) MyOne Streptavidin C1 beads (Invitrogen)). The DNA fragments were ligated with paired-end sequencing adaptors (Illumina, San Diego, CA) and then amplified (12-15 cycles). After amplification, the library was size-selected by agarose gel electrophoresis to remove DNA fragments whose length was less than 350 bp or more than 500 bp. The DNA fragments of 350-500 bp in the agarose gel were purified with a QIAquick gel extraction kit (QIAGEN). The size and concentration of the purified library were determined with a 2100 Bioanalyzer.

### Processing of the sequencing output

Image analysis and base calling were performed using Illumina RTA and CASAVA with a default parameter. Mapping and filtering were carried out according to a procedure reported by Imakaev *et al*. [5]. In brief, sequenced reads were mapped to the human reference genome (hg19) using Bowtie2 [6]. The reads of 25 bp were mapped at first. If the reads were not mapped uniquely, they were extended to 30 bp and re-mapped. This extension and re-mapping process was repeated until the read length of 75bp was reached. To remove non-informative pairs, the following read pairs were removed: (1) self circles (both reads mapped in the same restriction fragment), (2) dangling ends (sum of the distances between read and corresponding *Hin*dIII recognition site more than 500 bp), and (3) redundant (which may result from PCR amplification). We used a library “hiclib” (provided by the Leonid Mirny laboratory (https://bitbucket.org/mirnylab/)) for these filtering steps. The processing steps described above were carried out with our in-house computational pipeline.

### Expression quantitative trait loci (eQTL)

The GTEx Portal database provides SNPs associated with gene expression from 449 individuals not restricted to specific diseases or conditions in 44 diverse postmortem tissues. RNA expression was measured by RNA sequencing. Genotyping was performed on the Illumina Human Omni 2.5M and 5M Beadchips.

The BRAINAC database provides SNPs associated with gene expression from 134 neuropathologically normal individuals in 10 postmortem brain regions: cerebellar cortex, frontal cortex, hippocampus, inferior olivary nucleus (sub-dissected from the medulla), occipital cortex, putamen (at the level of the anterior commissure), substantia nigra, temporal cortex, thalamus (at the level of the lateral geniculate nucleus), and intralobular white matter. The BRAINAC database was calculated eQTL effects in the 10 brain regions and the average across all available regions. RNA expression was measured using an Affymetrix Exon 1.0 ST array. Genotyping was performed on the Illumina Infinium Omni1-Quad BeadChip.

### Publicly available gene expression datasets

syn5550404 includes gene expression data for 159 cerebellum (CBE) and 160 temporal cortex (TC) samples from North American Caucasian subjects with neuropathological diagnosis of AD (n=82) or elderly controls without neurodegenerative diseases (n=77 in CBE; n=78 in TC). All subjects were from the Mayo Clinic Brain Bank (MCBB) or the Banner Sun Health Research Institute. All ADs had definite diagnosis according to the the National Institute of Neurological and Communicative Disorders and Stroke and the Alzheimer’s Disease and Related Disorders Association (NINCDS-ADRDA) criteria and had Braak neurofibrillary tangle (NFT) stage of IV or greater. Control subjects had Braak NFT stage of III or less, the Consortium to Establish a Registry for Alzheimer’s Disease (CERAD) neuritic and cortical plaque densities of 0 (none) or 1 (sparse) and lacked any of the following pathologic diagnoses: AD, Parkinson’s disease, dementia with Lewy bodies, vascular dementia, progressive supranuclea palsy, motor neuron disease, corticobasal degeneration, Pick’s disease, Huntington’s disease, frontotemporal lobar degeneration, hippocampal sclerosis or dementia lacking distinctive histology. Gene expression measures were generated, using next-generation RNA sequencing with Illumina HiSeq 2000 sequencers.

In GSE5281, brain samples were collected at three Alzheimer’s Disease Centers (Washington University, Duke University, and Sun Health Research Institute). Individuals clinically classified as neurologically normal (10 males and 4 females) with a mean age of 79.8 ± 9.1 yr. Clinically classified late-onset AD-afflicted individuals (15 men and 18 women) with a mean age at death of 79.9 ± 6.9 yr. Samples were collected (mean postmortem interval (PMI) of 2.5 h) from six brain regions that are either histopathologically or metabolically relevant to AD and aging; these include the entorhinal cortex (BA 28 and 34), superior frontal gyrus (BA 10 and 11 and approximate BA 8), hippocampus, primary visual cortex (BA 17), middle temporal gyrus (BA 21 and 37 and approximate BA 22), and the posterior cingulate cortex (BA 23 and 31). Each brain tissues were analyzed on an Affymetrix Human Genome U133 Plus 2.0 Array.

In GSE44770, 230 autopsied tissues from dorsolateral prefrontal cortex (PFC), visual cortex (VC) and cerebellum (CR) in brains of LOAD patients, and non-demented healthy controls, collected through the Harvard Brain Tissue Resource Center (HBTRC), were profiled on a custom-made Agilent 44K array. All subjects were diagnosed at intake and each brain underwent extensive LOAD-related pathology examination. Gene expression analyses were adjusted for age and sex, PMI in hours, sample pH and RNA integrity number (RIN). In the overall cohort of LOAD and non-demented brains the mean ± SD for sample PMI, pH and RIN were 17.8±8.3, 6.4±0.3 and 6.8±0.8, respectively.

## References

1. Gatz M, Pedersen NL, Berg S, Johansson B, Johansson K, Mortimer JA, et al. Heritability for Alzheimer’s disease: the study of dementia in Swedish twins. J Gerontol A Biol Sci Med Sci. 1997;52:M117-25.

2. Harold D, Abraham R, Hollingworth P, Sims R, Gerrish A, Hamshere ML, et al. Genome-wide association study identifies variants at CLU and PICALM associated with Alzheimer’s disease. Nat Genet. 2009;41:1088–93.

3. Lambert J, Heath S, Even G, Campion D, Sleegers K, Hiltunen M, et al. Genome-wide association study identifies variants at CLU and CR1 associated with Alzheimer’ s disease. Nat Genet. 2009;41:1094–9.

4. Seshadri S, Fitzpatrick AL, Ikram MA, DeStefano AL, Gudnason V, Boada M, et al. Genome-wide analysis of genetic loci associated with Alzheimer disease. JAMA. 2010 12;303:1832–40.

5. Naj AC, Jun G, Beecham GW, Wang L-S, Vardarajan BN, Buros J, et al. Common variants at MS4A4/MS4A6E, CD2AP, CD33 and EPHA1 are associated with late-onset Alzheimer’s disease. Nat Genet. 2011;43436–41.

6. Lee JH, Cheng R, Barral S, Reitz C, Medrano M, Lantigua R, et al. Identification of Novel Loci for Alzheimer Disease and Replication of CLU, PICALM, and BIN1 in Caribbean Hispanic Individuals. Arch Neurol. 2011;68320–8.

7. Hollingworth P, Harold D, Sims R, Gerrish A, Lambert J-C, Carrasquillo MM, et al. Common variants at ABCA7, MS4A6A/MS4A4E, EPHA1, CD33 and CD2AP are associated with Alzheimer’s disease. Nat Genet. 2011;43429–35.

8. Lambert JC, Ibrahim-Verbaas CA, Harold D, Naj AC, Sims R, Bellenguez C, et al. Meta-analysis of 74,046 individuals identifies 11 new susceptibility loci for Alzheimer’s disease. Nat Genet. 2013;451452–8.

9. Schaub MA, Boyle AP, Kundaje A, Batzoglou S, Snyder M. Linking disease associations with regulatory information in the human genome. Genome Res. 2012;221748–59.

10. Kauwe JSK, Cruchaga C, Karch CM, Sadler B, Lee M, Mayo K, et al. Fine Mapping of Genetic Variants in BIN1, CLU, CR1 and PICALM for Association with Cerebrospinal Fluid Biomarkers for Alzheimer’s Disease. Bush A, editor. PLoS One. 2011;6e15918.

11. Soldner F, Stelzer Y, Shivalila CS, Abraham BJ, Latourelle JC, Barrasa MI, et al. Parkinson-associated risk variant in distal enhancer of α-synuclein modulates target gene expression. Nature. 2016;533:95–9.

12. Rosenthal SL, Barmada MM, Wang X, Demirci FY, Kamboh MI. Connecting the Dots: Potential of Data Integration to Identify Regulatory SNPs in Late-Onset Alzheimer’s Disease GWAS Findings. Arendt T, editor. PLoS One. 2014;9e95152.

13. Karch CM, Ezerskiy LA, Bertelsen S, Goate AM. Alzheimer’s Disease Risk Polymorphisms Regulate Gene Expression in the ZCWPW1 and the CELF1 Loci. Huang Q, editor. PLoS One. 2016;11e0148717.

14. GTEx Consortium. Human genomics. The Genotype-Tissue Expression (GTEx) pilot analysis: multitissue gene regulation in humans. Science. 2015;348:648–60.

15. French JD, Ghoussaini M, Edwards SL, Meyer KB, Michailidou K, Ahmed S, et al. Functional variants at the 11q13 risk locus for breast cancer regulate cyclin D1 expression through long-range enhancers. Am J Hum Genet. 2013;92:489–503.

16. Roussos P, Mitchell AC, Voloudakis G, Fullard JF, Pothula VM, Tsang J, et al. A role for noncoding variation in schizophrenia. Cell Rep. 2014;9:1417–29.

17. Tang Z, Luo OJ, Li X, Zheng M, Zhu JJ, Szalaj P, et al. CTCF-Mediated Human Genome Architecture Reveals Chromatin Topology for Transcription. Cell. 2015;163:1611–2.

18. Claussnitzer M, Hui C-C, Kellis M. FTO Obesity Variant and Adipocyte Browning in Humans. N Engl J Med. 2016;374:192–3.

19. ENCODE Project Consortium. An integrated encyclopedia of DNA elements in the human genome. Nature. 2012;48957–74.

20. Roadmap Epigenomics Consortium, Kundaje A, Meuleman W, Ernst J, Bilenky M, Yen A, et al. Integrative analysis of 111 reference human epigenomes. Nature. 2015;518317–30.

21. Ernst J, Kellis M. ChromHMM: automating chromatin-state discovery and characterization. Nat Methods. 2012;9215–6.

22. Ernst J, Kellis M. Large-scale imputation of epigenomic datasets for systematic annotation of diverse human tissues. Nat Biotechnol. 2015;33364–76.

23. Welter D, MacArthur J, Morales J, Burdett T, Hall P, Junkins H, et al. The NHGRI GWAS Catalog, a curated resource of SNP-trait associations. Nucleic Acids Res. 2014;42:D1001-6.

24. Lonsdale J, Thomas J, Salvatore M, Phillips R, Lo E, Shad S, et al. The Genotype-Tissue Expression (GTEx) project. Nat Genet. 2013;45:580–5.

25. Aguet F, Brown AA, Castel S, Davis JR, He Y, Jo B, et al. Genetic effects on gene expression across human tissues. Nature. 2017;550(7675):204–13.

26. Ramasamy A, Trabzuni D, Guelfi S, Varghese V, Smith C, Walker R, et al. Genetic variability in the regulation of gene expression in ten regions of the human brain. Nat Neurosci. 2014;171418–28.

27. International Genomics of Alzheimer’s Disease Consortium (IGAP). Convergent genetic and expression data implicate immunity in Alzheimer’s disease. Alzheimers Dement. 2015;11(6):658–71.

28. AMPAD Knowledge Portal Mayo Clinic RNAseq https://www.synapse.org/Portal.html#!Synapse:syn5550404

29. Liang WS, Reiman EM, Valla J, Dunckley T, Beach TG, Grover A, t al. Alzheimer’s disease is associated witheducedxpression ofnergy metabolism genes in posterior cingulateeurons. Proc Natl Acad Sci U S A. 2008;105:4441–6.

30. Zhang B, Gaiteri C, Bodea L-G, Wang Z, McElwee J, Podtelezhnikov AA, t al. ntegrated systems approach identifies geneticodes andetworks in late-onset Alzheimer’s disease. Cell. 2013;153:707–20.

31. Braak H, Braak E. Neuropathological stageing of Alzheimer-related changes. Acta Neuropathol. 1991;82:239–59.

32. Shih SJ, Allan C, Grehan S, Tse E, Moran C, Taylor JM. Duplicated downstream enhancers control expression of the human apolipoprotein E gene in macrophages and adipose tissue. J Biol Chem. 2000;275:31567–7.

33. Bekris LM, Lutz F, Yu C-E. Functional analysis of APOE locus genetic variation implicates regional enhancers in the regulation of both TOMM40 and APOE. J Hum Genet. 2012;57:18–2.

34. Dixon JR, Selvaraj S, Yue F, Kim A, Li Y, Shen Y, et al. Topological domains in mammalian genomes identified by analysis of chromatin interactions. Nature. 2012;485376–80.

35. Nora EP, Lajoie BR, Schulz EG, Giorgetti L, Okamoto I, Servant N, et al. Spatial partitioning of the regulatory landscape of the X-inactivation centre. Nature. 2012;485381–5.

36. Mifsud B, Tavares-Cadete F, Young AN, Sugar R, Schoenfelder S, Ferreira L, et al. Mapping long-range promoter contacts in human cells with high-resolution capture Hi-C. Nat Genet. 2015;47:598–606.

37. Won H, De La Torre-Ubieta L, Stein JL, Parikshak NN, Huang J, Opland CK, et al. Chromosome conformation elucidates regulatory relationships in developing human brain. Nature. 2016;538:523–7.

38. Kalhor R, Tjong H, Jayathilaka N, Alber F, Chen L. Genome architectures revealed by tethered chromosome conformation capture and population-based modeling. Nat Biotechnol. 2011;3090–8.

39. Schmid MW, Grob S, Grossniklaus U. HiCdat: a fast and easy-to-use Hi-C data analysis tool. BMC Bioinformatics. 2015;1–6.

40. Levy-Leduc C, Delattre M, Mary-Huard T, Robin S. Two-dimensional segmentation for analyzing Hi-C data. Bioinformatics. 2014;30i386–92.

41. Wendt KS, Yoshida K, Itoh T, Bando M, Koch B, Schirghuber E, et al. Cohesin mediates transcriptional insulation by CCCTC-binding factor. Nature. 2008;451:796–801.

42. Sexton T, Bantignies F, Cavalli G. Genomic interactions: chromatin loops and gene meeting points in transcriptional regulation. Semin Cell Dev Biol. 2009;20:849–55.

43. Nichols MH, Corces VG. A CTCF Code for 3D Genome Architecture. Cell. 2015;162:703–5.

44. Tang Z, Luo OJ, Li X, Zheng M, Zhu JJ, Szalaj P, et al. CTCF-Mediated Human 3D Genome Architecture Reveals Chromatin Topology for Transcription. Cell. 2015;163:1611–27.

45. Williams RL, Starmer J, Mugford JW, Calabrese JM, Mieczkowski P, Yee D, et al. fourSig: a method for determining chromosomal interactions in 4C-Seq data. Nucleic Acids Res. 2014;42:e68.

46. Wang Y, Zhang B, Zhang L, An L, Xu J, Li D, et al. The 3D Genome Browser: a web-based browser for visualizing 3D genome organization and long-range chromatin interactions. bioRxiv 112268; doi: https://doi.org/10.1101/112268.

47. Han H, Cho JW, Lee S, Yun A, Kim H, Bae D, et al. TRRUST v2: an expanded reference database of human and mouse transcriptional regulatory interactions. Nucleic Acids Res. 2018;4;46(D1):D380-D386.

48. Yokoyama K, Tezuka T, Kotani M, Nakazawa T, Hoshina N, Shimoda Y, et al. NYAP: a phosphoprotein family that links PI3K to WAVE1 signalling in neurons. EMBO J. 2011;30:4739–54.

49. Kunkle BW, Grenier-Boley B, Sims R, Bis JC, Naj AC, Boland A, et al. Meta-analysis of genetic association with diagnosed Alzheimer’s disease identifies novel risk loci and implicates Abeta, Tau, immunity and lipid processing. bioRxiv 294629; doi: https://doi.org/10.1101/294629.

50. Suzuki R, Lee K, Jing E, Biddinger SB, McDonald JG, Montine TJ, et al. Diabetes and insulin in regulation of brain cholesterol metabolism. Cell Metab. 2010;12:567–7.

51. Craft S, Watson GS. Insulin and neurodegenerative disease: shared and specific mechanisms. Lancet Neurol. 2004;3:169–7.

52. Barbero-Camps E, Fernández A, Martínez L, Fernández-Checa JC, Colell A. APP/PS1 mice overexpressing SREBP-2 exhibit combined Aβ accumulation and tau pathology underlying Alzheimer’s disease. Hum Mol Genet. 2013;22:3460–7.

53. Dixon JR, Selvaraj S, Yue F, Kim A, Li Y, Shen Y, et al. Topological domains in mammalian genomes identified by analysis of chromatin interactions. Nature. 2012;485376–80.

54. Nora EP, Lajoie BR, Schulz EG, Giorgetti L, Okamoto I, Servant N, et al. Spatial partitioning of the regulatory landscape of the X-inactivation centre. Nature. 2012;485381–5.

55. Rao SSP, Huntley MH, Durand NC, Stamenova EK, Bochkov ID, Robinson JT, et al. A map of the human genome at kilobase resolution reveals principles of chromatin looping. Cell. 2014;159:1665–8.

56. Ong C-T, Corces VG. CTCF: an architectural protein bridging genome topology and function. Nat Rev Genet. 2014;15:234–4.

57. Gallagher MD, Posavi M, Huang P, Unger TL, Berlyand Y, Gruenewald AL, et al. A Dementia-Associated Risk Variant near TMEM106B Alters Chromatin Architecture and Gene Expression. Am J Hum Genet. 2017;101:643–6.

58. Kaminski WE, Orsó E, Diederich W, Klucken J, Drobnik W, Schmitz G. Identification of a novel human sterol-sensitive ATP-binding cassette transporter (ABCA7). Biochem Biophys Res Commun. 2000;273:532–8.

59. Fu Y, Hsiao J-HT, Paxinos G, Halliday GM, Kim WS. ABCA7 Mediates Phagocytic Clearance of Amyloid-β in the Brain. J Alzheimers Dis. 2016;54:569–84.

60. Pei B, Sisu C, Frankish A, Howald C, Habegger L, Mu XJ,t al. The GENCODE pseudogeneesource. Genome Biol. 2012;13:R51.

61. Storey JD, Tibshirani R. Statistical significance for genomewide studies. Proc Natl Acad Sci U S A. 2003;100:9440–5.

62. Tripathi S, Pohl MO, Zhou Y, Rodriguez-Frandsen A, Wang G, Stein DA, t al. Meta- and Orthogonalntegration ofnfluenza “OMICs” Data Defines a Role for UBR4 in Virus Budding. Cell Host Microbe. 2015;18:723–35.

63. Imakaev M, Fudenberg G, McCord RP, Naumova N, Goloborodko A, Lajoie BR, et al. Iterative correction of Hi-C data reveals hallmarks of chromosome organization. Nat Methods. 2012;9:999–1003.

## Supplementary References

1. Kalhor, R., Tjong, H., Jayathilaka, N., Alber, F. and Chen, L. (2012) Genome architectures revealed by tethered chromosome conformation capture and population-based modeling. Nature Biotechnology, 30, 90-98.

2. Liu, J. and Francke, U. (2006) Identification of cis-regulatory elements for MECP2 expression. Human Molecular Genetics, 15, 1769-1782.

3. Murrell, A., Heeson, S. and Reik, W. (2004) Interaction between differentially methylated regions partitions the imprinted genes Igf2 and H19 into parent-specific chromatin loops. Nature Genetics, 36, 889-893.

4. Lieberman-Aiden, E., van Berkum, N.L., Williams, L., Imakaev, M., Ragoczy, T., Telling, A., Amit, I., Lajoie, B.R., Sabo, P.J., Dorschner, M.O. et al. (2009) Comprehensive mapping of long-range interactions reveals folding principles of the human genome. Science, 326, 289-293.

5. Imakaev, M., Fudenberg, G., McCord, R.P., Naumova, N., Goloborodko, A., Lajoie, B.R., Dekker, J. and Mirny, L.A. (2012) Iterative correction of Hi-C data reveals hallmarks of chromosome organization. Nature Methods, 9, 999-1003.

6. Langmead, B. and Salzberg, S.L. (2012) Fast gapped-read alignment with Bowtie 2. Nature Methods, 9, 357-359.

